# Transient Kinetic Proofreading

**DOI:** 10.1101/2021.03.10.434642

**Authors:** G. Iyengar, M. Perry

## Abstract

We propose a new stochastic model for understanding the transient kinetic proofreading mechanism in a T-cell. Our model indicates that a stochastic version of absolute ligand discrimination is a consequence of the finite number of receptors on the cell surface; thus, pointing to receptor number control as being critical to T-cell activation. We propose four different metrics to characterize the performance of kinetic proofreading mechanisms. We explore the numerical experiments that explore the trade-offs between speed, specificity, sensitivity, and robustness of T-cell activation as a function of the model parameters. We also consider the impact of receptor clustering on these trade-offs.

## 1 Introduction

Kinetic proofreading is a mechanism that explains the extremely low probability of error observed in a number of biological processes, e.g. microtubule growth, DNA replication, and the activation of T-cells by antigen presenting cells (***Hopfield, 1974***; ***Ninio, 1975***). Kinetic proofreading exploits the fact that cycling through intermediate complexes on the path to a final product can ensure that the path to undesired products has a significantly lower probability than the path to the desired final product. In this work, we are concerned with kinetic proofreading in switching systems where the cell takes one of two actions based on environmental signals. In particular, we will focus on T-cell activation.

T-cells are a component of the adaptive arm of the immune system (***Perelson and Weisbuch, 1997***). In contrast to the innate immune system, which attacks a broad range of pathogens based on common cell-surface markers, each individual cell in the adaptive immune system is specially tailored to sense and react to antigens associated with a specific type of pathogen. Among other things, a T-cell is responsible for sensing and killing pathogens expressing the T-cell’s target antigen on their cell surface. T-cells are able to sense very small differences in the binding energy between a specific “non-self” antigen and T-cell receptors (TCRs) and that of a “self” antigen and the TCR to activate only in the presence of the “non-self” antigen. Kinetic proofreading has been proposed as a mechanism that allows T-cells to discriminate with much better accuracy than that implied by the difference in binding energy alone.

***McKeithan (1995***) proposed the first kinetic proofreading-based model for TCR activation. This model is described by the following set of ODEs ***François and Altan-Bonnet (2016***).

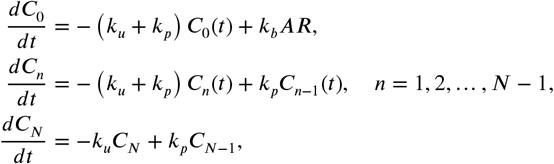

where *A* denotes the density of the antigen interacting with the TCR, *R* denotes the density of the TCR, *C*_0_(*t*) is the density of a TCR-antigen complex at time *t, C*_*n*_(*t*) is the density of a TCR-antigen complex with *n* phosphorylations at time *t, k*_*b*_ is the rate at which antigens bind to TCRs and is assumed to be the same for all antigens, *k*_*u*_ is the rate at which antigens unbind from TCRs and varies between different antigens, and *k*_*p*_ is the rate at which TCR-antigen bindings are phosphorylated. The affinity *τ* of an antigen is defined as the reciprocal of the unbinding rate *k*_*u*_, i.e. *τ* = 1/*k*_*u*_. The model assumes that a TCR-antigen complex with *N* phosphorylations is required for T-cell activation.

The steady state value of *C*_*N*_ ∝ *L*/(*k*_*u*_)^*N*^. Suppose a self antigen with affinity 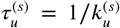 and a non-self antigen with affinity 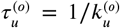 are present with the same concentration *R*. Then the ratio of the steady-state concentration 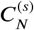 of the self-antigen-TCR complex and the steady-state concentration 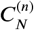 of the non-self-antigen-TCR complex is 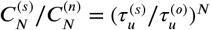. Thus, by repeatedly dissociating the receptor-antigen bindings en route to a final product, the proofreading network ensures that the rate of error decreases exponentially with the size of the network.

The analysis above assumes that the concentration of self and non-self antigens are equal. This is almost never the case. There is evidence of densities as low as ten non-self antigens per cell leading to T-cell activation (***Irvine et al., 2002***), whereas self antigens are far more abundant. T-cells activate even to non-self antigen at very low concentrations, and do not activate even when self antigen is present in very high concentrations. The performance of a kinetic proofreading mechanism can be summarized by its *activation set* 𝒜 which is defined as the set of pairs (*L, τ*) for which the mechanism activates.

The ability of the T-cells to distinguish between self and non-self antigens despite very large discrepancies in concentration is known as *absolute ligand discrimination* (***François and Altan-Bonnet, 2016*). *François and Altan-Bonnet (2016***) (see, also (***Altan-Bonnet and Germain, 2005*; *Lalanne and François, 2013*))** proposed and analyzed deterministic mean-field models for absolute ligand discrimination that were based on a feedback loop that regulates the phosphorylation rate of antigen-bound TCRs via a kinase whose activity is itself regulated by the quantity of phosphorylated antigen-bound TCRs. This feedback loop, called *adaptive sorting*, decouples the behavior of the proofreading mechanism from antigen density, thereby achieving absolute ligand discrimination. Although the impact of this feedback loop on T-cell ligand discrimination in vivo is questionable (***Paster et al., 2015***), decoupling the behavior of the proofreading network from antigen affinity is key for absolute discrimination. The activation set 𝒜^*I*^ corresponding to *ideal* absolute ligand discrimination is given by 𝒜^*I*^ = {(*L, τ*) : *L* ≥ *L*_min_, *τ* ≥ *τ*_*min*_}, i.e. the lower boundary of the set 𝒜 is given by an asymptote in the *L*-direction located at *τ*_min_ and an asymptote in the *τ*-direction located at *L*_min_. See Figure 1.

**Figure 1.**
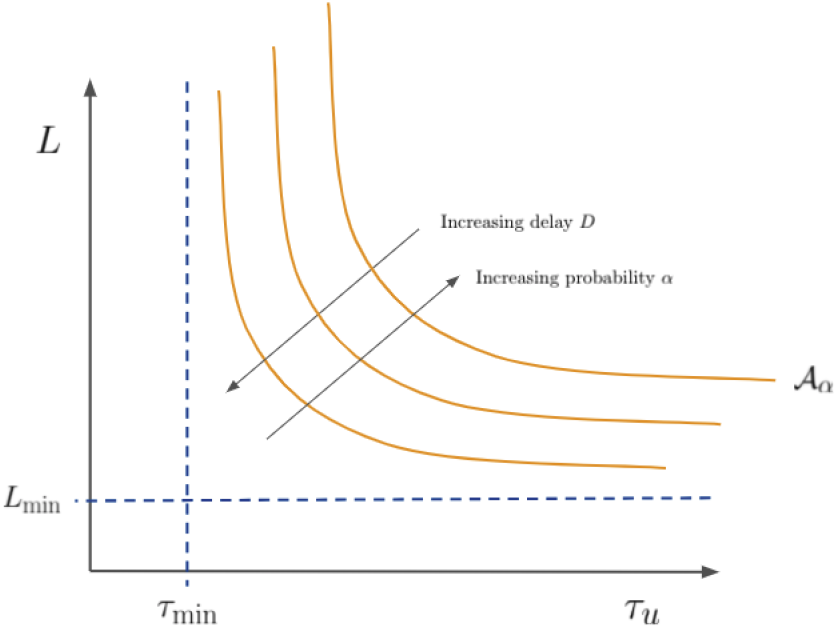
The activation set 𝒜_*α*_ bounded by the vertical asymptote at *τ*_min_ and the horizontal asymptote at *L*_min_ defining the ideal absolute ligand discrimination activation set 𝒜^*I*^. The delay *D* can be fixed and the probability *α* of activation increased to move 𝒜_*α*_ up and to the right, and the probability of activation can be fixed and the delay can be increased to move 𝒜_*α*_ down and to the left.

The existing models of absolute ligand distribution are both mean field and steady state. Therefore, these models are not able to represent the trade-off between the delay and other performance measures. Furthermore, these approaches are also not able to model the impact of stochastic noise. In this work, we propose a model that accounts for both transience and stochasticity. We assume the kinetic proofreading mechanism needs to make the activation decision within a specified delay *D*, and therefore the decision inherently involves stochastic analysis. The performance metric of interest is now the probability *α* ∈ (0, 1) by which the mechanism activates by delay *D*. Consequently, in the stochastic setting, the activation set 𝒜_*α*_ is now indexed by a probability *α*. Following standard practice, we set *α* = 0.95. See Figure 1.

The focus of the existing work on kinetic proofreading is to develop mechanisms that yield error-correction, and absolute ligand discrimination models have focused on the existence of asymptotes in the *L* and *τ* directions. We propose that these mechanisms must be evaluated on several other dimensions. In particular, we propose the following four metrics:

a. *Sensitivity*: The level *L*_*α*_ of the asymptote in the *τ*_*u*_-dimension, i.e. the lowest density at which the cell activates within delay *D* with probability at least *α* for any value of the affinity *τ*_*u*_.
b. *Speciicity*: The gap *τ*_0.95_ − *τ*_0.05_ between the asymptotes along the *L*-direction for *α* = 0.05 and *α* = 0.95, i.e. the difference between the affinity values for which the cell almost never activates (i.e. *α* = 0.05) and almost always activates (i.e. *α* = 0.95) in the limit of large density. The affinity of the non-self antigen should be greater than *τ*_0.95_, and the affinity of the self-antigen should be less than *τ*_0.05_ to guarantee that the mechanism makes very few errors. Therefore, the smaller the gap, the more specific is the response of the mechanism.
c. *Fidelity*: This metric quantifies the distance of the activation set 𝒜_*α*_ associated with a mechanism, and that associated with ideal absolute ligand discrimination, i.e. the distance of the curve from the two asymptotes. We provide one possible definition of this metric in Section 2.3.
d. *Robustness*: This metric measures how the activation set changes when the parameters defining the mechanism are perturbed. Robustness is an important metric since noise becomes very important at low copy numbers.

Although versions of these metrics apply to any proofreading mechanism involved in switching, we trade-off between these desirable properties in the context of a particular model for absolute ligand discrimination.

We propose a new stochastic model where a T-cell activates when the total number of antigen-TCR-downstream complexes exceeds a certain threshold *Z*. We show how to efficiently compute the probability that the T-cell activates with a certain delay *D*, allowing us to investigate the impact of delay on each performance metric.

In our model absolute ligand discrimination emerges as a direct consequence of the fact that the number of TCRs on the cell surface is finite. This connection between absolute ligand discrimination and receptor number control (***Sorkin and Von Zastrow, 2009***) opens up the possibility that the properties of a single T-cell can be modulated by controlling the number of receptors.

T-cell activation in our model is controlled by the number of TCRs *N*, the threshold *Z* on the number of antigen-TCR-downstream molecule complexes, the delay *D*, and the probability *α*. We show how these parameters trade-off sensitivity, specificity, fidelity, and robustness. We also show how to use the model to explore the impact of receptor clustering, and show that such clustering improves fidelity and robustness to biochemical errors.

The rest of this paper is organized as follows. In Section 2.1 we introduce the model, and in Section 2.2 show that the model achieves absolute ligand discrimination because the number of receptors on the cell surface is finite. In Section 2.3 we discuss the results of numerical experiments investigating the trade-offs between sensitivity, specificity, fidelity, speed, and robustness. In Section 2.4 we discuss the impact of clustering on ligand discrimination, and in Section 3 we conclude with a discussion of possible experiments to test the different behaviors seen in the model.

## 2 Results

### 2.1 Model

Consider a single T-cell exposed to a single antigen with a uniform density *L* on the cell surface. The cell has *N* receptors spread uniformly across its surface that bind to the antigen. We work with the following approximation for the T-cell activation dynamics. An antigen-bound receptor can recruit one more molecule that we refer to as Lck ^1^, and the total number of antigen-TCR-Lck complexes needs to be larger than a threshold for the T-cell to activate. It will be clear from the discussion below, one could add more downstream molecules to the model without changing the analysis or the computation.

Following ***Husain (2018***); ***Taylor et al. (2017***), we model the response of a T-cell to an antigen by a two-dimensional stochastic model. The events in the model are as follows:

1. A single T-cell has a finite number *N* of TCRs available on its surface. The assumption that *N* is finite is a significant departure from ***Husain (2018***); ***Taylor et al. (2017***), and we show that this is directly leads to absolute ligand discrimination.
2. An unbound TCR binds to an antigen with rate *k*_*b*_*L*. We implicitly assume that the TCR only reacts to a single antigen, and are ignoring the effects of ligand antagonism (***François et al., 2016***).
3. Antigen-bound TCRs recruit a single Lck molecule at rate *k*_*on*_.
4. Antigens unbind from TCRs at rate 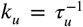. We assume that the antigen-TCR-Lck complex loses the Lck molecule if the antigen unbinds. In line with ***McKeithan (1995***), we assume that Lck’s do not unbind from antigen-TCR complexes unless the complex itself breaks apart. This assumption does not change the qualitative behavior of the model.
5. The cell activates once *Z*(≥2) TCRs have recruited Lck molecules ^2^.

The state of a T-cell in our model is described by two variables: the number of antigen-TCR complexes *n* ≤ *N* and the number of antigen-TCR-Lck complexes *z* ≤ *Z*. We let (*n*_*t*_, *z*_*t*_) denote the state of the T-cell at any time *t* ≥ 0. Since *n*_*t*_, *z*_*t*_ ≥ 0, *z*_*t*_ ≤ *n*_*t*_, and the cell activates when *z*_*t*_ = *Z*, it follows that the state space is

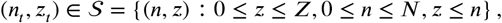

and the dynamics of {(*n*_*t*_, *z*_*t*_) : *t* ≥ 0} is described by a two-dimensional continuous time Markov chain with an absorbing boundary at

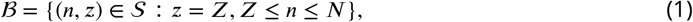

i.e. ℬ refers to all cell states corresponding to an activated cell. See Figure 2 for a schematic of the Markov chain transitions and Appendix A for more details on the model. In our numerical simulations, we use 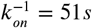 and 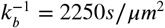 (***Husain, 2018***; ***Taylor et al., 2017***).

**Figure 2.**
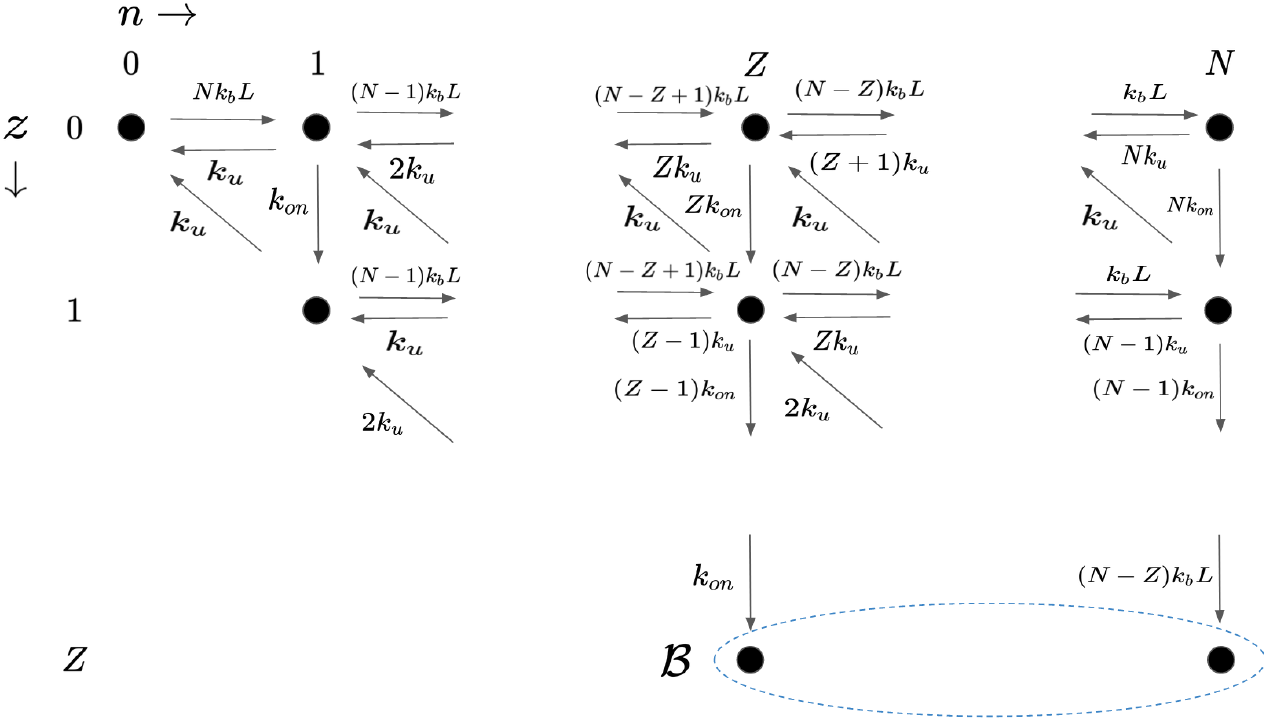
State space and transition rates for model of T-cell activation. Horizontal movements correspond to changes in the number of antigen-bound TCRs *n*, while vertical movements correspond to changes in the number of antigen-TCR-Lck complexes *z*. ℬ corresponds to states for which the cell activates.

We assume that the T-cell must make a decision within a specified delay *D* from the first presentation of the antigen, i.e. if the cell does not have *Z* Lck’s recruited by time *D*, the cell has missed its opportunity to sense the antigen in time. This is a departure from previous kinetic proofreading models, in which activation is a function of the steady-state concentrations and information theoretic tools can be used to analyze the system (***Cover, 1999***; ***Ganti et al., 2020***; ***Das et al., 2018***). We want to characterize the set 𝒜_*α*_ of density-affinity pairs (*L, τ*_*u*_) for which the T-cell activates before *D* with probability *α* ∈ (0, 1). In order to characterize the operating characteristics of the T-cell, we define the first passage time *T*_(*i,j*)_ from the state (*i, j*) ∈ 𝒮\ℬ to a state (*n, Z*) ∈ ℬ as follows:

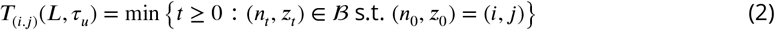

i.e. *T*_(*i,j*)_ is the first time at which a cell, initially with *i* ≥ 0 antigen-bound TCRs and 0 ≤*J* ≤ *i* antigen-TCR-Lck complexes, has *Z* antigen-TCR-Lck complexes. We emphasize that *T*_(*i,j*)_ is a function of the density-affinity pair (*L, τ*_*u*_) since the rate matrix *Q* (see Appendix for details) depends on (*L, τ*_*u*_). Since the random time to activation after the initial presentation of the antigen is given by *T*_(0,0)_(*L, τ*_*u*_), it follows that

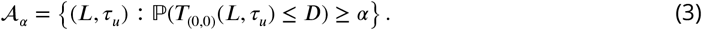

In order to characterize 𝒜_*α*_, we need to characterize the entire distribution of *T*_(0,0)_(*L, τ*_*u*_) as opposed to just the mean first passage time (MPFT), as was done in earlier works (***Husain, 2018***; ***Taylor et al., 2017***). We are interested in characterizing sets 𝒜_*α*_ for a range of *α* in order to understand the impact of the probability threshold *α*.

Since an analytical expression for the distribution of the first passage time *T*_(0,0)_ is unknown, we compute and invert the Laplace transform 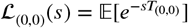 of *T*_(0,0)_ to compute the entire cumulative distribution function (CDF) ℙ (*T*_(0,0)_ ≤*t*) for all *t* ≥ 0. We show that the complexity of computing the entire distribution is comparable to computing the MPFT. Furthermore, since we compute the entire CDF, we can compute the expected value of other performance measures. See Appendix A for details.

### 2.2 Finite number of receptor leads to absolute ligand discrimination

Let

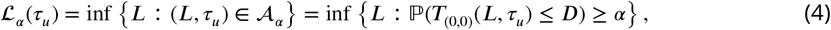

denote the lowest density for an antigen with affinity *τ*_*u*_ that ensures that the cell activates within a delay *D*. In Figure 3 we plot ℒ_*α*_ (*τ*_*u*_) vs *τ* for *α* ∈ {0.05, 0.5, 0.95}. The set 𝒜_*α*_ is above and to the right of the corresponding ℒ_*α*_ -curve. We also plot the minimum density required for the mean first passage time (MFPT) to be below the delay *D*. The plots in this figure also clearly show the additional information in the collection of plots ℒ_*α*_ for different values of *α*. For example, the plots show that the set 𝒜_*α*_ is much more sensitive to affinity *τ*_*u*_ as compared to the density *L*. This information is not available in the single plot corresponding to the MFPT.

**Figure 3.**
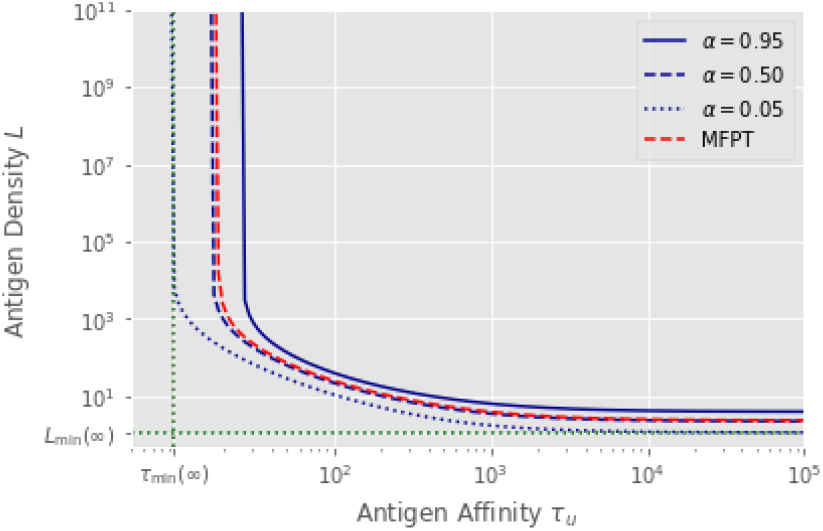
Let ℒ_*α*_ (*τ*_*u*_) denote the minimum density required to activate by delay *D* with probability *α* (see (4)). Plots of ℒ_*α*_ (*τ*_*u*_) for *Z* = 8 *N* = 12 and delay *D* = 10^3^. For each affinity *τ*_*u*_, the minimum density *L* was found using bisection over densities ranging from 10^−4^ to 10^16^. The delay *D* = 10^3^ seconds was chosen because it represents a realistic amount of time for human immune cells to operate over (10^3^ seconds ≈ 15 minutes). Horizontal and vertical asymptotes *L*_min_ and *τ*_min_ displayed for *α* = 0.05. Also included is the minimum density *L* needed for a cell exposed to antigen with affinity *τ*_*u*_ to have a MFPT less than *D*. Time to compute *α* = 0.50 curve was 30 seconds, time to compute MFPT curve was 2 seconds. This factor of 15 difference comes from using 15 summands to approximate the Bromwich integral. This same ratio holds when testing larger *N* and *Z* values (see section A in the appendix).

The ℒ_*α*_ curve is very closely approximated by two asymptotes: an asymptote in the *L*-direction at the affinity *τ*_min_, and an asymptote in the *τ*_*u*_-direction at the density *L*_min_. (See Figure 3 for the asymptotes corresponding to the ℒ_0.05_ (*τ*_*u*_) curve.) The thresholds (*L*_min_, *τ*_min_) are functions of the parameters of the mechanism (*N, Z*), the allowed delay *D*, and the probability threshold *α*. This asymptotic behavior directly leads to a probabilistic version of the absolute ligand discrimination described in ***Lalanne and François (2013***). For antigens with sufficiently large affinity, the cell activates with very high probability by the cutoff time *D*, and it does so for all antigen density *L* above some threshold *L*_min_. For a fixed density above this lower threshold, one can decrease the affinity until a critical affinity *τ*_min_ is reached, at which point the probability of activation starts to dramatically decrease as the affinity is further decreased. Finally, this critical affinity is the same for almost all densities above the density threshold.

We show that the asymptotes are an immediate consequence of the fact that *N* is finite, and do *not* exist if *N* is allowed to be infinite. First consider the asymptote in the *L*-direction. For large enough densities *L*, all of the *N* TCRs are very quickly bound to an antigen, and therefore the Lck dynamics, and hence the activation probability, are independent of density *L*. We rigorously prove this in section C in the appendix.

The dynamics that lead to the asymptote in the *τ*_*u*_-direction are more subtle. Justifying the existence of a strictly positive density *L*_min_ below which the T-cell never activates for any value of the affinity *τ*_*u*_ is equivalent to showing that 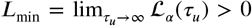. Since increasing the affinity only makes it easier for the TCR-antigen-Lck complex to recruit and hold on to Lck, it is clear that ℒ_*α*_ (*τ*_*u*_) is a decreasing function of *τ*_*u*_. So *L*_min_ ≥ ℒ_*α*_ (∞). When *τ*_*u*_ = ∞, antigens never never unbind, and so both *n*_*t*_ and *z*_*t*_ are non-decreasing over time. Since *z*_*t*_ ≤ *n*_*t*_, it follows that ℙ (*z*_*t*_ ≥ *Z*) ≤ ℙ (*n*_*t*_ ≥ *Z*) i.e. the likelihood of the cell being in an activated state by time *t* must be less than the likelihood of the cell having *Z* antigen-bound TCRs at time *t*. Furthermore, since the rate of recruiting one more antigen-bound TCR is bounded above by *Nk*_*b*_*L*, it follows that the probability that there are *Z* antigen-bound TCRs by time *D* is less than 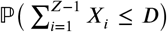, where each *X*_*i*_ is independent, exponentially distributed with rate *Nk*_*b*_*L*. Thus, it follows that 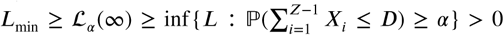 for sufficiently large *D* (see section B in the appendix). Note that the last term is strictly positive only if *N* < ∞ i.e. we lose the existence of the non-zero horizontal asymptote when the number of receptors is infinite. We rigorously establish this result in section D in the appendix.

We conclude by noting that absolute ligand discrimination holds even in the presence of additional downstream molecules. Suppose that each antigen-bound TCR must recruit *M* > 1 additional molecules, each at rate *k*_*on*_, and that the cell activates once it has *Z*_*M*_ TCRs that have recruited *M* molecules. The ℒ_*α*_ curves for this model will still have a vertical asymptote i.e. some minimum affinity *τ*_min_ below which no antigen will activate the cell in time with sufficient probability, even if the antigen has extremely high density. This occurs for the same reason mentioned before - for sufficiently high antigen density *L*, all *N* TCRs are bound to an antigen with very high probability for all points in time. Therefore, the dynamics of the proofreading network in this limit is effectively independent of the affinity *τ*_*u*_. The ℒ_*α*_ curves will also have a horizontal asymptote i.e. a density *L*_min_ below which no antigen will activate the cell in time even with extremely high affinity, and this holds due to the exact same argument above that the time until activation is lower-bounded by the time to recruit *Z*_*M*_ antigens. These arguments, however, do not explain how properties of the kinetic proofreading network scale as the dimension of the model is increased. We leave this discussion for future work.

### 2.3 Sensitivity, specificity, fidelity, and robustness

In the Introduction, we introduced four metrics that we argued effectively characterize the performance of a kinetic proofreading mechanism: specificity, sensitivity, fidelity and robustness. In this section, we define these metrics for our proposed mechanism. In order to define these metrics, we need to define the following function:

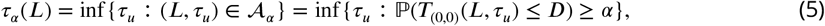

i.e. *τ*_*α*_ (*L*) is the minimum affinity at which the proofreading network activates with probability *α* by delay *D* when the antigen density is *L*. Note that *τ*_min_ = *τ*_*α*_ (∞) is the *x*-axis location of the asymptote in the *L*-direction. Next, we define the four metrics.

a. *Speciicity*: The level of the horizontal asymptote ℒ_0.95_(∞) for *α* = 0.95. Clearly, the smaller the sensitivity, the better the network is at responding to small amounts of antigen.
b. *Speciicity*: The relative gap (*τ*_0.95_(∞) − *τ*_0.05_(∞))/ *τ*_0.95_(∞) between the *x*-axis location of the asymptote in the *L*-direction for *α* = 0.95 and *α* = 0.05. High specificity implies that the likelihood of activation decreases quickly as the antigen affinity is decreased. Therefore, networks with higher specificity are better at discriminating between two affinity values.
c. *Fidelity*: The first two metrics refer to the asymptotes. For the ideal proofreading network, the ℒ_*α*_ curve is the *L*-shaped curve defined by the two asymptotes. Fidelity is a metric that measures how far ℒ_*α*_ is from the ideal *L*-shape. We need to define some intermediate quantities in order to define fidelity. Fix *ϵ* > 0. We say that the curve ℒ_*α*_ (*τ*) reaches within a factor (1 + *ϵ*) of the asymptote in the *τ*_*u*_-direction at *τ*_*ϵ*_ = *τ*_*α*_((1 + *ϵ*)*L*_min_), and reaches within a factor (1 + *ϵ*) of the asymptote in the *L*-direction at *L*_*ϵ*_ = ℒ_*α*_ ((1 + *ϵ*) *τ*_min_). See Figure 4. Monotonicity properties of the model imply that the ℒ_*α*_ (*τ*_*u*_) curve lies in the following two *L*-shaped regions:

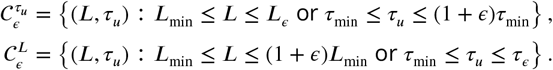 The set 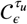 would represent an ideal proof-reading curve provided the maximum of the *x*-width *ϵτ*_min_ and *y*-width (*L*_*ϵ*_ − *L*_min_) is close to zero, i.e. 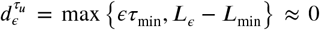. Similarly, define 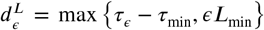), and 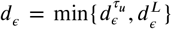 to be the minimum of the distance to ideal performance implied by either of the two covering sets. We define the *fidelity F* (*N, Z, D, α*) as the inverse of the minimum “distance” over all possible choices for *ϵ*, i.e.

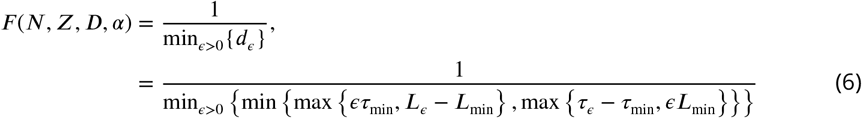 Note that high fidelity implies that the transition is closer to ideal behavior.
d. *Robustness*: Since the actual number of TCRs *N* + *δN* on the cell surface may differ from *N* because of transport noise, the corresponding ℒ_*α*_ curve may no longer be able to discriminate between the self and non-self antigens. We define the robustness *r*_*N*_ (*N, Z, D*) of the kinetic proofreading network with parameters (*N, Z, D*) to errors in *N* as follows:

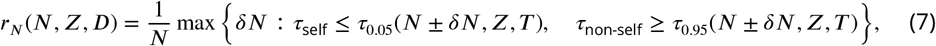 i.e. *r*_*N*_ (*N, Z, D, α*) is the maximum allowed relative change in *N* before the kinetic proofreading network is unable to differentiate between the self and non-self antigens. The robustness in *Z* is defined as

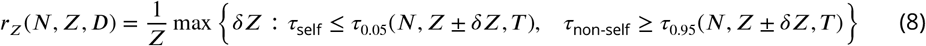 It is possible that the robustness *r*_*N*_ (*N, Z, D, α*) is very sensitive to *z*. In order to study this effect, we define robustness *r*_*NZ*_(*N, Z, D, α*) to simultaneous perturbations in *N* and *z* as follows:

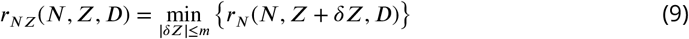

**Figure 4.**
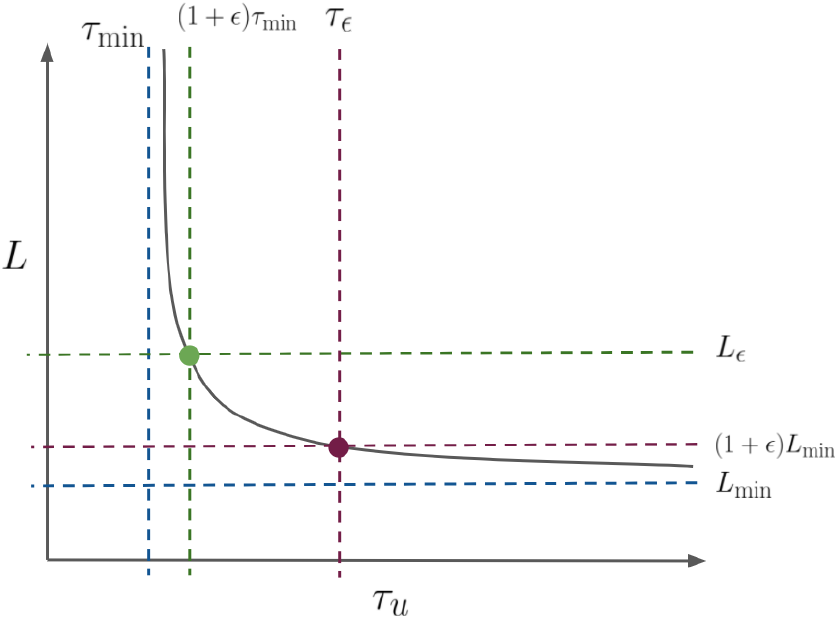
Examples of covering (*τ*_*u*_, ℒ_*α*_ (*τ*_*u*_)) curve with 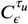 (in green) and 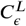(in red).

To summarize, a kinetic proofreading network that exhibits perfect absolute ligand discrimination has zero specificity, zero sensitivity, and infinite fidelity. Next, we investigate the possible values of sensitivity, specificity, and fidelity achievable by the proposed Markov chain model as a function of the parmeters (*N, Z, D*).

We simulated the performance of the kinetic proofreading system for

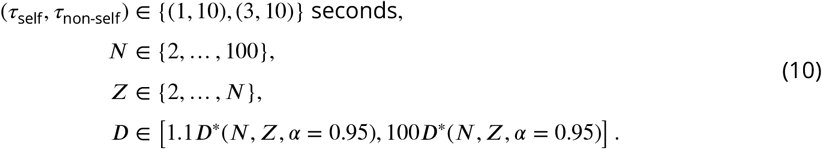

We call (*N, Z, D*) *admissible* for the pair of self and non-self antigen affinities (*τ*_self_, *τ*_non-self_) if the *α* = 0.95 vertical asymptote *τ*_0.95_(*N, Z, D*) ≤ *τ*_non-self_, and the *α* = 0.05 vertical asymptote *τ*_0.05_(*N, Z, D*) ≥ *τ*_self_. This filtering step fixes the specificity of the proofreading network.

In Figure 5 we plot sensitivity *L*_min_(*N, Z, D, α* = 0.95) vs the delay *D* for all admissible values of (*N, Z, D*) – each point corresponds to an admissible triple (*N, Z, D*). The lower envelop of the set of points corresponds to values of (*N, Z, D*) that are Pareto optimal with respect to sensitivity and delay, i.e. these points represent the optimal trade-off between these two metrics. Note that on the Pareto front, the sensitivity *L*_min_ is a decreasing function of the delay *D*, i.e. the proofreading network can be designed to be more sensitive if a higher delay is tolerable.

**Figure 5.**
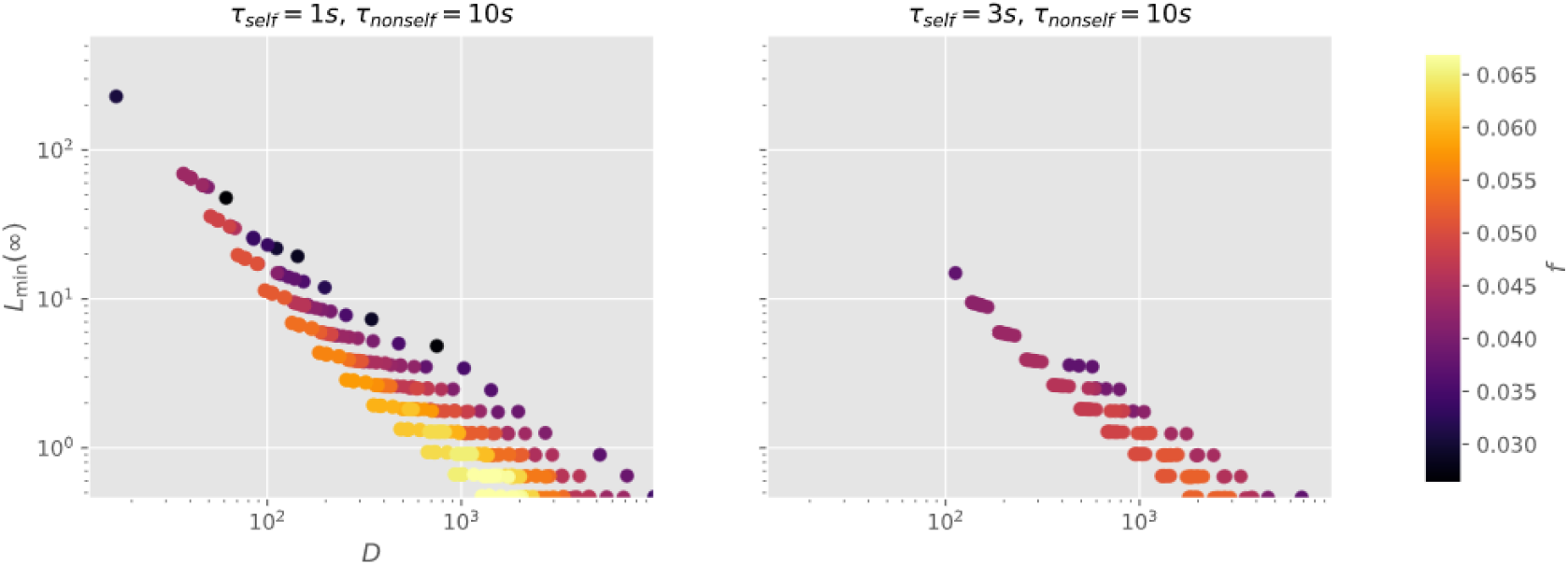
Achievable values of *D* and *L*_min_ for fixed *τ*_self_ and *τ*_non-self_. Points are colored based on the value of *F* (*L, D*).

Next, we want to visualize the (*L, D, F*) Pareto front. There are often several close together (*L, D*) values that result in very different values for fidelity. In order to account for this, we take the maximum value of fidelity in a neighborhood of (*L, D*), and define

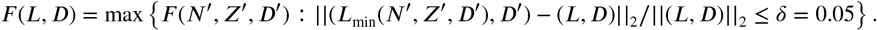

The color of the point (*L, D*) in figure 5 denotes the value of *F* (*L, D*). On the Pareto front, the fidelity and sensitivity both increase as the allowed delay *D* increases, and begins to level off between *D* = 100 and *D* = 1000 seconds, a biologically realistic range of activation times. In the right-hand plot in Figure 5 we see that, as one would expect, increasing the required specificity by increasing *τ*_self_ limits the delay, sensitivity, and fidelity with which the cell can operate.

In Figure 6 the points are colored by robustness. In the first row the points are colored by robustness to perturbations in *N*, i.e. *r*_*N*_ defined in (7), in the second row the points are colored by robustness to perturbations in *z*, i.e. *r*_*Z*_ defined in (8), and the third row the points are colored by the joint robustness *r*_*NZ*_ defined in (9). We see that the values of *r*_*N*_ and *r*_*Z*_ on the Pareto front increase as *D* increases. The trends in the *N* and *z*-robustness *r*_*NZ*_ are slightly more complicated – on the Pareto front, *r*_*NZ*_ is maximized for intermediate values of *D*. Increasing specificity by only a small amount (e.g. a 30% increase in *τ*_self_) can significantly reduce the achievable level of robustness in the kinetic proofreading model. Increasing specificity had the same effect on fidelity, although this change is not as drastic as that for robustness.

**Figure 6.**
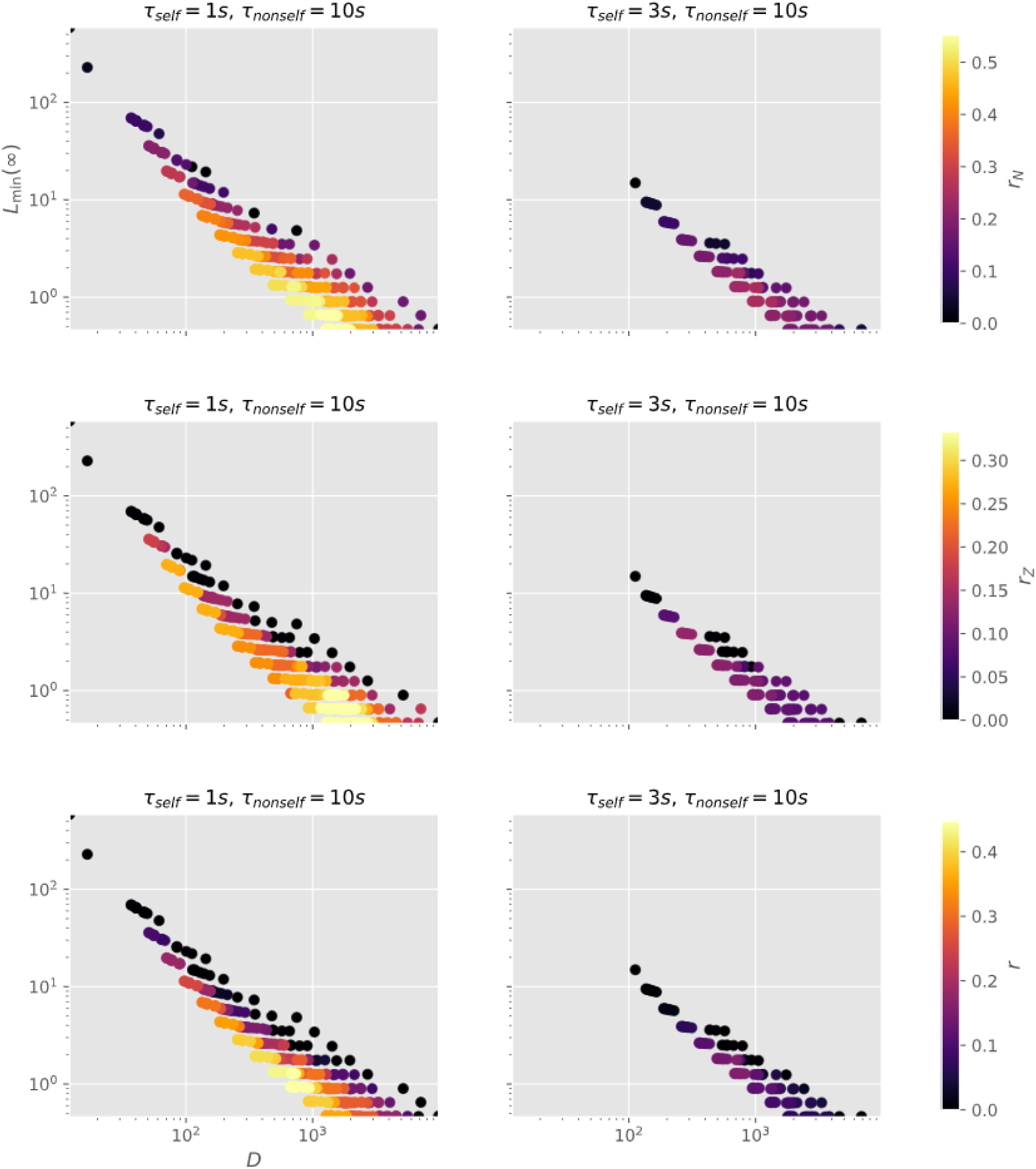
Achievable values of *D* and *L*_min_ for fixed *τ*_self_ and *τ*_non-self_. Points in the top two plots are colored based on *r*_*N*_. Points in the middle two plots are colored based on *r*_*Z*_. Points in the bottom two plots are colored based on *r*_*NZ*_. In all three rows, highest robustness among nearby points is used to color the points.

### 2.4 Clustering

***Husain (2018***); ***Taylor et al. (2017***) proposed the model that we refine in this work to understand the role of receptor clustering in the activation of T-cells. Spatial heterogeneity and clustering of cell surface receptors is of general interest as well (***Burroughs and Van Der Merwe, 2007***; ***Iyengar and Rao, 2014***). ***Taylor et al. (2017***) found that clustering of antigen-bound TCRs is crucial for ligand discrimination in T-cells. We consider here what a cell potentially gains by incorporating clustering into its proofreading mechanism.

To approximate the impact of clustering we modify the original model as follows. We assume that the cell surface has two compartments, each initially having *N* TCRs. The dynamics of the two compartments are assumed to be independent. We let *L*_*i*_ denote the antigen density on compartment *i* = 1, 2, and say that the cell activates once one of the compartments has recruited *Z* Lck molecules. We introduce volatility in the antigen density by setting *L*_1_ = (1 − *ϵ*)*L* and *L*_2_ = (1 + *ϵ*)*L* for some *ϵ* ∼ uniform(−1, 1), so that the average antigen density over the whole surface is still *L*. We assume that the cell is able to sense density and places (1 + *δ*)*N* receptors in the compartment where the density is higher, and (1 − *δ*)*N* receptors in compartment with the lower density.

Relaxing the assumptions of this simple model i.e. that the antigen density on each compartment is fixed over time, the cell surface only has two compartments, the cell can pre-emptively move *δN* receptors from compartment 1 to compartment 2, and that the number of receptors in each compartment stays fixed over time, is left for future analysis.

Figure 7 plots admissible (*L*_min_, *D*) values at a specified level of specificity for models without clustering or variation in antigen density, without clustering but with variation in density, with clustering but without variation in density, and with both both clustering and variation in density. Points are colored based on fidelity. Figure 8 is the same as figure 7 but with points colored by robustness to simultaneous errors in *N* and *Z*.

**Figure 7.**
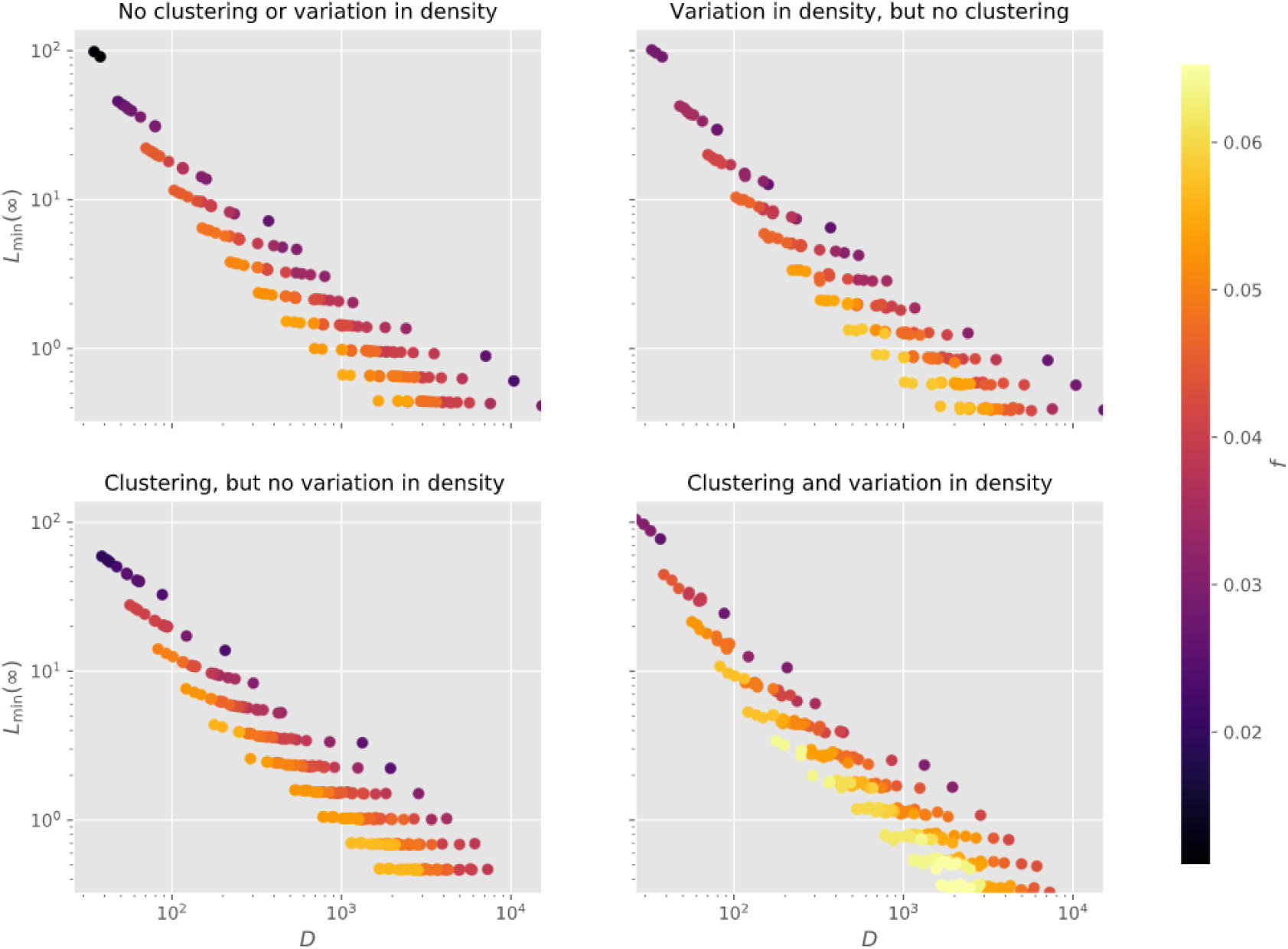
Achievable values of *D* and *L*_min_ for *τ*_self_ = 1s and *τ*_nonself_ = 10s. Points colored based on fidelity. Plots in top row have no clustering (*δ* = 0.0), plots in bottom row have clustering with *δ* = 0.50. Plots in left column have no variation in antigen density (*ϵ* = 0.0), whereas plots in right column have variation in density with *ϵ* ∼ uniform(−1, 1). For plots with variation in density, probability of activation was calculated with 20 samples of *ϵ*.

**Figure 8.**
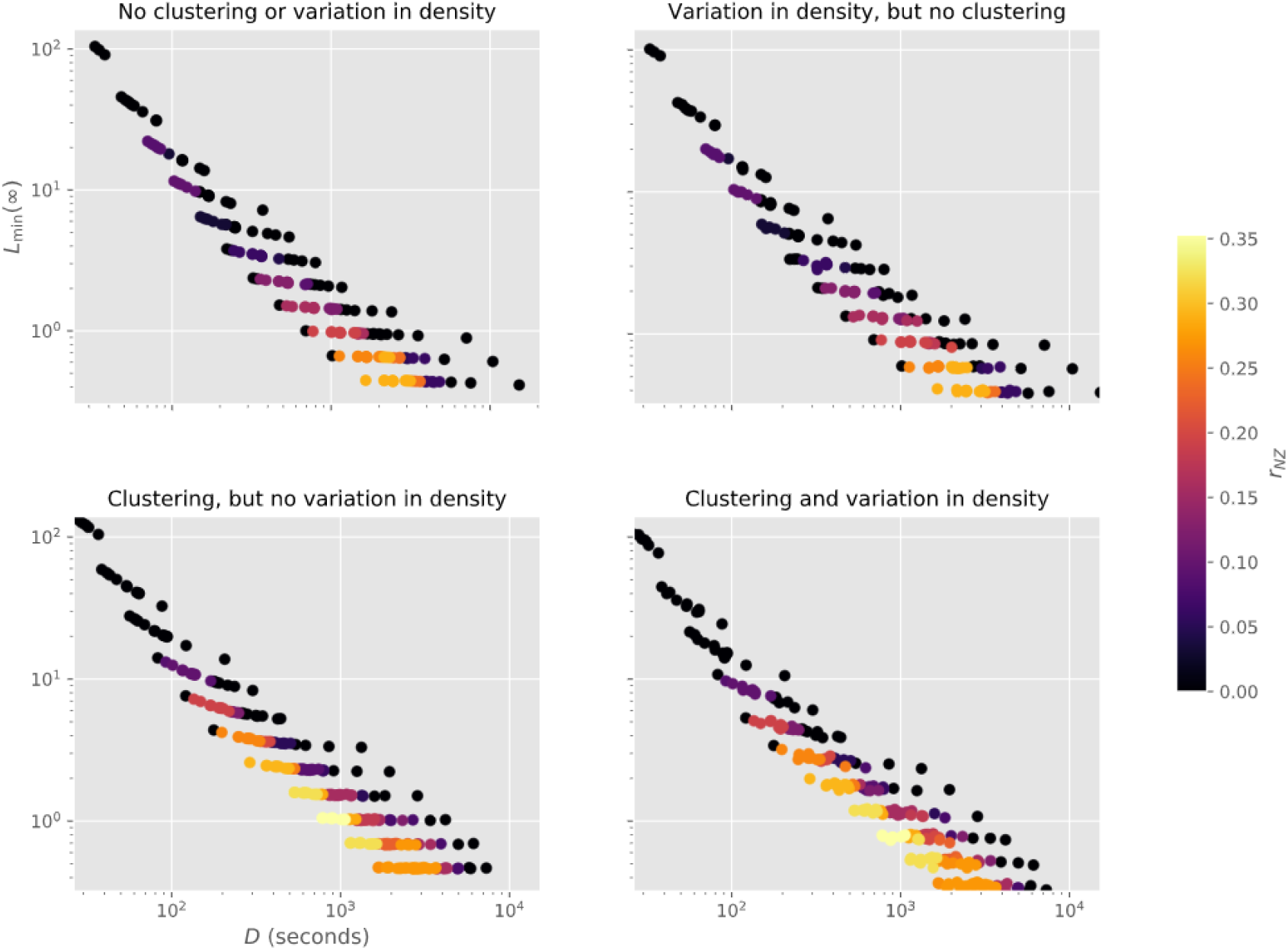
Same layout and parameters as figure 7, but with points colored based on robustness to simultaneous errors in *N* and *Z*.

From figure 7 we see that clustering enhances the fidelity of the proofreading network given the existence of variation in the antigen density, with no loss in sensitivity or speed at the given level of specificity. A similar result is seen in figure 8, with clustering improving the robustness to simultaneous errors in the number of surface receptors *N* and activation threshold *Z*, albeit regardless of whether or not there’s variation in the antigen density. The impact of clustering on robustness to *N* and *Z* separately is similar, and figures for these quantities are omitted. These figures point towards clustering as a potential way of improving the fidelity and robustness of proofreading.

## 3 Discussion

We propose a kinetic proofreading model that exhibits a stochastic version of absolute ligand discrimination as a direct consequence of controlling the number of receptors. We propose that the performance of a kinetic proofreading mechanism can be characterized by sensitivity, specificity, fidelity, robustness to biochemical errors and delay. We explored the trade-off between these different metrics within our proposed model. We also examined the impact of receptor clustering on these trade-offs.

In Figure 9 we plot the predictions of our model as the number of receptors *N* is changed. Our model predicts a decrease in the level *L*_min_ of the horizontal asymptote as *N* increases; furthermore, the rate of decrease in *L*_min_ is decreasing in *N*, i.e. successive doubling of *N* does not result in equal amounts of decrease in *L*_min_. A similar prediction holds for *τ*_min_ as well. Moreover, the model predicts that the transition from the horizontal asymptote to the vertical one happens over a larger interval of affinity values as *N* increases. These predictions can be experimentally validated by inhibiting or enhancing receptor endocytosis (***Sorkin and Von Zastrow, 2009***) and observing the impact on antigen sensing.

**Figure 9.**
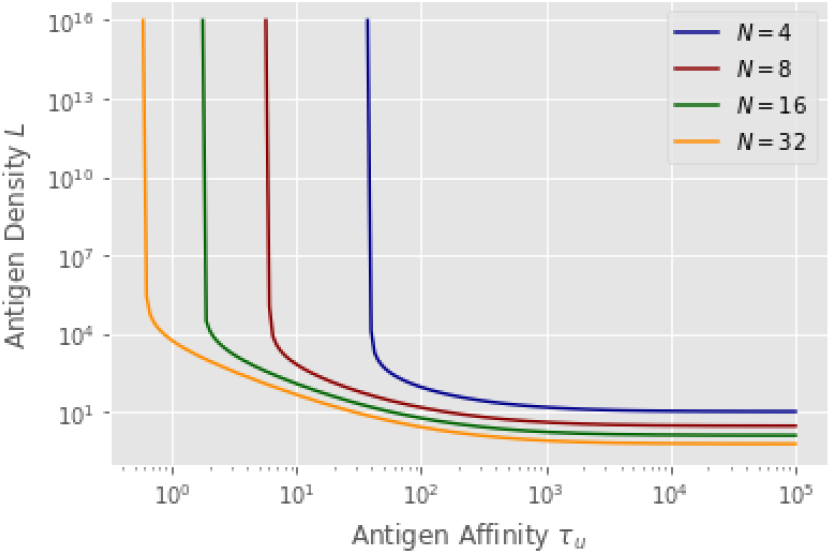
ℒ_*α*_ plots for various values of *N. Z* = 4, *α* = 0.95, *D* = 10^3^ seconds.

We also focused on the trade-offs between various metrics in the proposed kinetic proofreading model. Given the many different functions performed by cells and the various environments they must operate in, one would expect there to be differences in how cells prioritize sensitivity, specificity, speed, and robustness. Controlling the number of surface receptors, the threshold *Z* on the downstream molecule, and the delay *D*, may be a way for cells to trade-off these metrics. Given different priorities among cells, it could be worth surveying different types of T-cells, as well as B-cells ***Tsourkas et al. (2012***), and seeing whether the quantity of receptors displayed on their surface matches up with what they prioritize.

The model also showed that clustering may serve the purpose of increasing the fidelity and robustness of proofreading, particularly when the antigen density varies over the cell surface. It may be interesting to attempt an experiment where the extracellular fluid around a collection of T-cells is agitated so as to induce heterogeneities in antigen density. The model here would predict more clustering the more the fluid is perturbed.

As a final thought, our the model can be modified to study the effect of cell heterogeneity. Biological systems are subject to large amounts of noise that can both help and hurt the system. Developing this model further to study how cell heterogeneity can help a population of T-cells be more robust to changing environmental conditions ***Kussell and Leibler (2005***) and antigen variability ***Mayer et al. (2015***), and how the noisy responses from heterogeneous cells can be aggregated in an effective manner ***Butler et al. (2013***), could be promising extensions.

## A Model details

The full rate matrix *Q* for the model described in section 2.1 is defined as follows (***Sigman, 2009***).

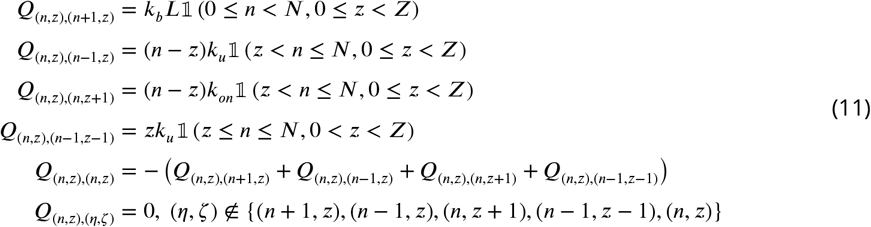

These rates follow from the assumptions described in the text. For example, if the T-cell is in state (*n, z*) and it has not yet activated, i.e. *z* < *Z*, the antigen binds to an available TCR, i.e. if *n* < *N*, with rate *k*_*b*_*L*; thus, *Q*_(*n,z*),(*n*+1,*z*)_ = *k*_*b*_*L* if *n* < *N* and *z* < *Z*. All the other rates similarly follow from the assumptions above. Note that *Q*_(*n,z*),(*n,z*)_ is set to ensure that ∑ _(*n*′,*z*′) ∈ 𝒮_ *Q*_(*n,z*),(*n*′,*z*′)_ = 0.

The first step in computing the distribution of *T*_(0,0)_ is to compute the Laplace transform ℒ_(0,0)_(*s*) of *T*_(0,0)_.

### Lemma A.1 (Laplace Transform)

*Let* 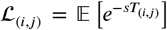 *denote the Laplace transform of the density function for the irst passage time T*_(*i,j*)_ *from state* (*i, j*) *to the absorbing boundary* ℬ. *Then*

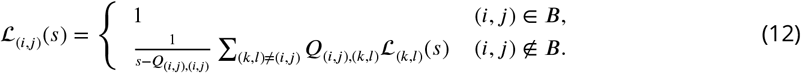

*Therefore*, ℒ_(0,0)_(*s*) *can be computed by solving the above set of equations. The complexity of computing* ℒ_(0,0)_ *is* 𝒪 (*N*^2^*Z*^2^).

*Proof.* Since ℬ is absorbing, it follows that *T*_(*i,j*)_ = 0 for all (*i*.*j*) ∈ ℬ. Therefore, ℒ _(*i,j*)_ (*s*) = 1 for all *s*. Fix (*i, j*) ∉ ℬ. Then the first-step analysis approach implies that ***Privault (2013***)

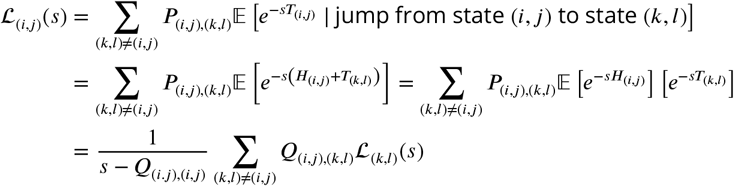

where 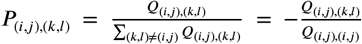 is the probability of transitioning to state (*k, l*) ≠ (*i, j*) upon leaving (*i, j*), and *H*_(*i,j*)_ is the holding time for state (*i, J*) and is exponentially distributed with parameter −*Q*_(*i,j*),(*i,j*)_.□

The cumulative density function (CDF) is given by

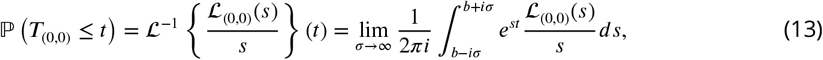

where *b* is any real number to the right of all singularities of ℒ_(0,0)_(*s*)/*s*. Let *s* = *c* + *id*, for *c, d* ∈ ℝ. Then

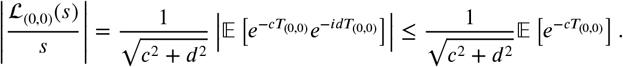

Thus, it follows that any singularity of ℒ_(0,0)_(*s*)/*s* has **ℜ**(*s*) ≤ 0, and therefore any *b* > 0 is sufficient. We use *b* = 1 in our numerical study. Next, using the trapezoidal rule with step size *h*, we approximate the Bromwich intergral in (13) with an infinite series ***Abate et al. (2000***)

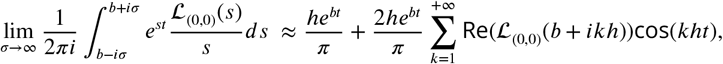

and appropriately truncate the series to approximate the integral. For *h* and the number of terms in the approximation, we use the specification in ***Abate et al. (2000***). We found that the number of terms was independent of the parameters (*N, Z, D*), i.e. only a constant number of Laplace transform evaluations are needed for any parameter values. Thus, it follows that the computational complexity of computing the CDF in terms of (*N, Z*) is the same as that of computing the mean first passage time (MFPT), since computing the MFPT reduces to solving a system of equations very similar to those used to compute ℒ_(0,0)_.

It is also clear that this method of computing the first passage time until activation will still work even if the number of downstream molecules that can bind to each TCR is increased to *M* > 1, albeit the system of equations to solve for the Laplace transforms 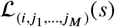 of the first passage times 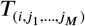 until activation (which in this case corresponds to the cell reaching some critical threshold *Z*_*M*_ antigen-bound TCRs with *M* additional molecules recruited) will take 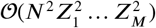 time.

## B Minimum possible activation time

Throughout the text we implicitly assume that the delay *D* is chosen to be large enough so that it’s possible for the cell to activate quickly enough with sufficiently high probability. In this section, we compute a lower bound for the delay *D* such that ℙ (*T*_(0,0)_(*L, τ*_*u*_) ≤ *D*) ≥ *α* for some (*L, τ*_*u*_).

Since the forward transitions, i.e. those that increase (*n, z*), are increasing in *L* and the backward transitions, i.e. those that decrease (*n, z*), are decreasing in *τ*_*u*_, it follows that the first passage time *T*_(0,0)_ is monotonically decreasing in both *L* and *τ*_*u*_. Thus, in order to ensure there exist (*L, τ*_*u*_) for which ℙ (*T*_(0,0)_(*L, τ*_*u*_) ≤*D*) ≥ *α*, it is sufficient to show that 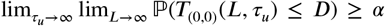. In Section C, we establish that the first passage time *T*_(0,0)_(*τ*_*u*_, *L*) in the limit *L* → ∞ is approximated by the first passage time 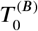 from state 0 to state *z* in a birth-death process with birth rates. *λ*_*z*_ = (*N* − *z*)*k*_*on*_ and death rate *µ*_*z*_ = *z*/*τ*_*u*_. Next, taking the limit *τ*_*u*_ → ∞, the limit

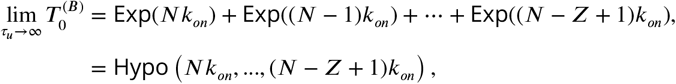

where Exp(*λ*) denotes an exponential random variable with rate *λ* and Hypo(*λ*_1_, …, *λ*_*k*_) is a hypo-exponential distribution defined by the given rates. The same limit results from taking the limit *L* → ∞ in the Markov chain *M*^(*f*)^ that we discuss in Section D. Thus, in order to ensure that the network reaches ℬ with probability at least *α*, we need

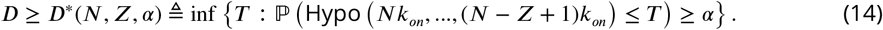

We ensure that *D* ≥ *D*^*^(*N, Z, α*) in all of our analyses so that *τ*_min_(∞) and *L*_min_(∞) are well-defined.

## C Vertical asymptote

Fix *D* ≥ *D*^*^ (see (14)), and define

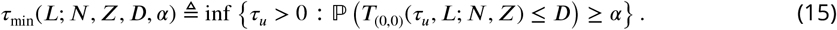

where *T*_(0,0)_(*τ*_*u*_, *L*; *N, Z*) is the first passage time from state (0, 0) to the absorbing boundary B for an antigen with affinity *τ*_*u*_ and density *L* when the cell has *N* TCRs and activates when *Z* antigen-TCR-Lck complexes form. Informally, *τ*_min_(*L*; *N, Z D, α*) denotes the minimum affinity *τ*_*u*_ needed to ensure that the cell activates with probability at least *α* by time *D* with antigen density *L*, when the cell has *N* TCRs on the cell surface, and *Z* antigen-TCR-Lck complexes are needed for activation. For brevity, let *τ*_min_(*L*) denote *τ*_min_(*L*; *N, Z, D, α*).

To study the behavior of *τ*_min_(*L*) as *L* → ∞, we define

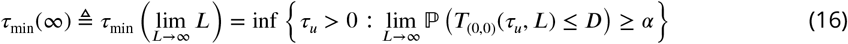

where the last equality holds by Fatou’s lemma and the fact that lim_*L*→ ∞_ {*T*_(0,0)_(*τ*_*u*_, *L*) ≤ *D*} exists and is equal to {*T*_(0,0)_(*τ*_*u*_, ∞) ≤ *D*}, so we can exchange the limit and probability.

Note that as *L* → ∞, all the *N* TCRs instantly convert into TCR-antigen complexes, and the bottle-neck event that determines activation is the recruitment of Lck, i.e. activation is controlled by the Lck dynamics. We formalize this in Theorem C.1. Let *B* = {*b*_*t*_ : *t* ≥ 0} denote a birth-death process defined on {*z* : 0 ≤ *z* ≤ *Z*} with birth rates *λ*_*z*_ = (*N* − *z*)*k*_*on*_ and death rates *µ*_*z*_ = *z*/*τ*_*u*_, and an absorbing boundary at *Z*. Let 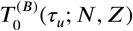 denote the first passage time from state 0 to state *Z* in a birth-death process *B*. Then, we have the following characterization for *τ*_min_(∞).

### Theorem C.1 (Vertical Asymptote)

*The minimum ainity τ*_min_(∞) *required to activate in the limit L* → ∞ *is given by*

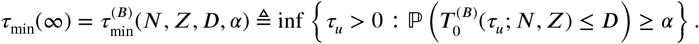

Thus, Theorem C.1 implies that, in order to characterize *τ*_min_(∞) as a function of (*N, Z, T, α*), we only need to compute the distribution of the first passage time 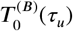 for the birth-death process *B*. The Laplace transforms ℒ _*i*_ (*s*) for the first passage time from state *i* to state *Z* for *B* can be computed in the same way as the Laplace transforms ℒ_(*i,j*)_(*s*) for the original model. Computing ℒ_*i*_(*s*) for 0 ≤ *i* ≤ *Z* reduces to solving a tri-diagonal system of equations, with a computational complexity of 𝒪(*Z*) as opposed to 𝒪(*N*^2^*Z*^2^). Figure D.1 confirms that 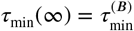 and that 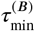 can be computed much faster than *τ*_min_(∞) (∼10-100 times faster for moderate-sized *N* and *Z*).

*Proof.* This result follows from showing that 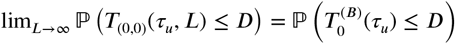. Let (*n*_*t*_, *z*_*t*_) be the state of the Markov chain at time *t*. Since the boundary ℬ is absorbing,

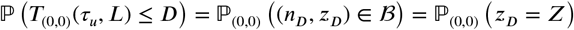

with ℙ_(*n,z*)_ denoting the distribution of (*n*_*t*_, *z*_*t*_) given (*n*_0_, *z*_0_) = (*n, z*). Therefore, it suffices to show 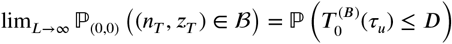. We establish this by showing

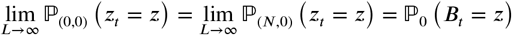

The first equality is established in Lemma C.2, and the second equality is established in Lemma C.3. □

The intuition here is that in the limit of *L* → ∞, all *N* TCRs are instantaneously bound to antigens, and the activation is determined by the dynamics of the antigen bound TCRs acquiring Lck molecules. In Lemma C.2 we rigorously establish the first result.

### Lemma C.2.

*For all t* > 0, lim_*L*→ ∞_ ℙ_(0,0)_(*z*_*t*_ = *z*) = lim_*L*→ ∞_ ℙ_(*N*,0)_(*z*_*t*_ = *z*).

*Proof.* The rate out of each state (*n, z*) in the chain is upper bounded by 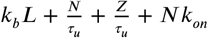. We define the following four independent Poisson processes.

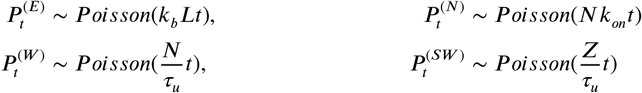

We think of the arrivals for the process 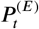 as “eastward” movements from (*n, z*) to (*n, z*+1), arrivals for 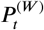 as “westward” movements from (*n, z*) to (*n* − 1, *z*), arrivals for 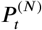 as “northward” movements from (*n, z*) to (*n, z* + 1), and arrivals for 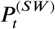 as “southwestward” movements from (*n, z*) to (*n* − 1, *z* − 1). We can see that

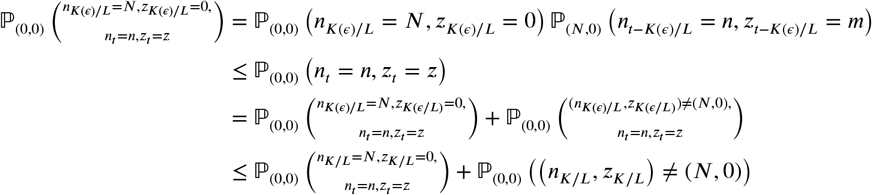

Next, we have

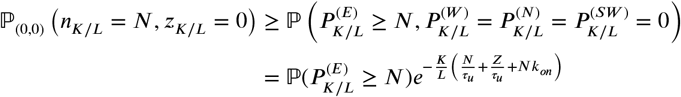

Since *e*^−*x*^/*x*^*j*^ → 0 as *x* → ∞ for any finite *j*, it follows that 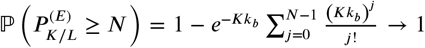 as *K* → ∞, and 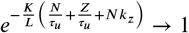 as *L* → ∞ for any fixed *K*. Thus, it follows that for all *ϵ* > 0, there exists some *K*= *K*(*ϵ*) such that for all *L* ≥ *L*(*ϵ*), 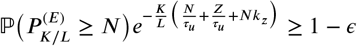. Hence,

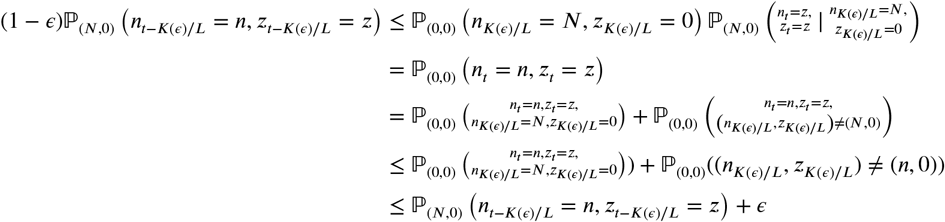

Now, choose any sequence of densities {*L*_*k*_} with *L*_*k*_ → ∞, as well as a sequence {*ϵ* _*k*_} such that *ϵ* _*k*_ → 0 and *K*(*ϵ* _*k*_)/*L*_*k*_ → 0. Taking the limit along (*L*_*k*_, *ϵ* _*k*_) gives

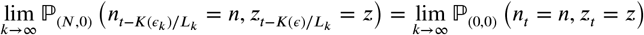

In the limit *k* → ∞, *K*(*ϵ* _*k*_)/*L*_*k*_ → 0, and the state distribution of a continuous time Markov chain is continuous in time, so

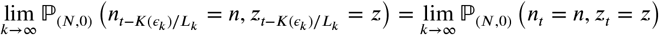

and we can conclude that lim_*L*→ ∞_ ℙ _(0,0)_ (*n*_*t*_ = *n, z*_*t*_ = *z*) = lim_*L*→ ∞_ ℙ _(*N*,0)_ *n*_*t*_ = *n, z*_*t*_ = *z*). □

Next, we establish that dynamics of the 2D Markov chain in the limit of *L* → ∞ is given by the 1D birth-death process in the Lck dimension.

### Lemma C.3.

lim_*L*→ ∞_ ℙ_(*N*,0)_(*z*_*t*_ = *z*) = ℙ_0_(*B*_*t*_ = *z*).

*Proof.* Instead of tracking the aggregate behavior (*n*_*t*_, *z*_*t*_) of the chain, we keep track of individual arrivals and departures of TCRs and Lcks, and show lim_*L*→ ∞_ ℙ _(*N*,0)_(*z*_*t*_ ≤ *z*) = ℙ_0_(*B*_*t*_ ≤ *z*) using the following coupling

1. Let (*a*_*t*_, *d*_*t*_) denote the birth and death processes associated with the birth-death process *B*_*t*_ i.e. *B*_*t*_ = *a*_*t*_ − *d*_*t*_.
2. Suppose there is an arrival at time *s*. Then, increase *z*_*s*_ by 1 with probability *n*_*s*_/*N* and keep it the same with probability 1 − *n*_*s*_/*N*.
3. Suppose there is a departure at time *s*, decrease *z*_*s*_ by 1 only if the associated arrival had led to an increase in *z*_*s*_.

By construction, *z*_*t*_ defined above has the same distribution as the number of Lcks *z*_*t*_ in the Markov chain. Also by construction, *z*_*t*_ ≤ *B*_*t*_ on a sample-path basis, and therefore ℙ_(*N*,0)_(*z*_*t*_ ≤ *z*) ≥ ℙ_0_(*B*_*t*_ ≤ *z*), i.e. lim_*L*→ ∞_ ℙ_(*N*,0)_(*z*_*t*_ ≤ *z*) ≥ ℙ_0_(*B*_*t*_ ≤ *z*).

Additionally, *B*_*t*_ − *z*_*t*_ ≤ *A*(*T*_*N*_ (*t*)) on a sample-path basis, with *T*_*N*_ (*t*) being the total amount of time the Markov chain spends in states with *n*_*s*_ < *N* for *s* ∈ (0, *t*], and *A*(*T*_*N*_ (*t*)) being the number of arrivals in *B*_*t*_ during the periods of time in (0, *t*] when *n*_*s*_ < *N*. Thus, we have that

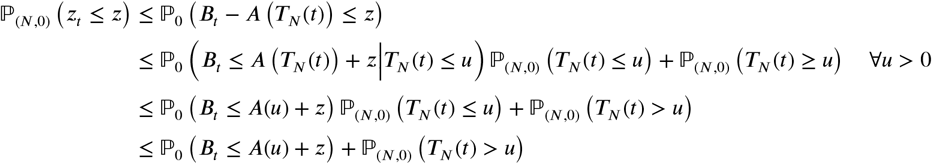

where the second-to-last inequality uses the fact that *T*_*N*_ (*t*) is independent of *B*_*t*_. Taking limits, we see

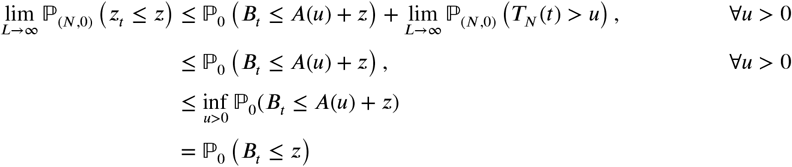

The first line uses the fact that *B*_*t*_ and *A*(*u*) are independent of *L*. The second inequality comes from the fact lim sup_*L*→ ∞_ ℙ_(*N*,·)_ (*T*_*N*_ (*t*) > *u*)= 0 for all *u* > 0, with ℙ_(*n*,·)_ being the conditional probability when the chain starts with *n* TCRs and arbitrary 0 ≤ *z* ≤ *n* at time 0. This result is established in Lemma C.4. The third inequality comes from the fact that, in general, the CDF of a random variable is monotonically increasing in its argument, and *A*(*u*) is monotonically decreasing in a sample-path sense, so ℙ_0_ (*B*_*t*_ ≤ *A*(*u*) + *z* is minimized by taking *u* → 0. The last equality uses the fact that CDFs by definition are right-continuous, and lim_*u*→ ∞_ *A*(*u*) = 0 in a sample-path sense. We conclude that lim_*L*→ ∞_ ℙ_(*N*,0)_ (*z*_*t*_ ≤ *z*) = ℙ_0_ (*B*_*t*_ ≤ *z*). □

### Lemma C.4

lim sup_*L*→ ∞_ ℙ_(*N*,·)_(*T*_*N*_ (*t*) ≥ *u*) = 0.

*Proof.* By Markov’s inequality, we know

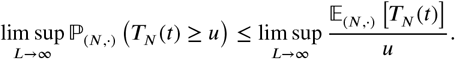

Next, let *D*_*N*_ (*t*) be the number of transitions from a state with *N* TCRs to a state with *N* − 1 TCRs in (0, *t*] and *S* be the sojourn time from a state with *N* − 1 TCRs to a state with *N* TCRs. By definition, 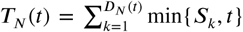 with the *S*_*k*_ ‘s being iid copies of *S*, so by Wald’s equation we have 𝔼_(*N*, ·)_[*T*_*N*_(*t*)] = 𝔼_(*N*, ·)_[*D*_*N*_(*t*)]𝔼_(*N*−1, ·)_[min{*S,t*}]. We see that

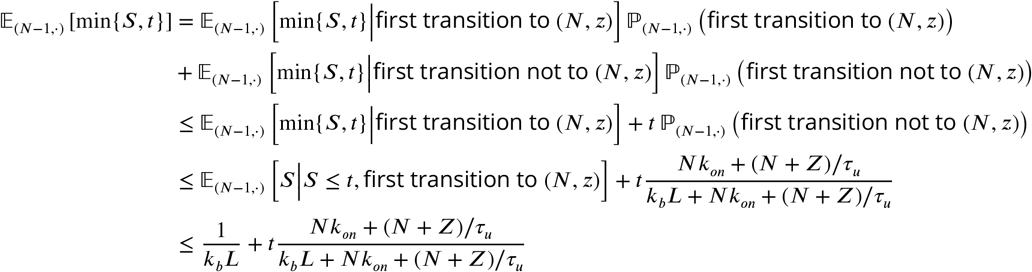

Since the rates for the Poisson processes 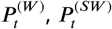, and 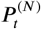 are all finite for all *τ*_*u*_ > 0 and independent of *L*, 𝔼_(*N*,·)_ [*D*_*N*_ (*t*)] is finite for all *L*. Therefore, we conclude that lim sup_*L*→ ∞_ 𝔼_(*N*,·)_ [*T*_*N*_ (*t*)] = 0, and lim sup_*L*→ ∞_ ℙ_(*N*,·)_ (*T*_*N*_ (*t*) ≥ *u*)= 0. □

## D Horizontal asymptote

Fix *D* ≥ *D*\* (see (14)), and let

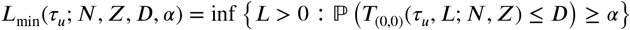

denote the minimum density *L* needed to ensure that the T-cell activates with delay *D* with probability at least *α*. For brevity, let *L*_min_(*τ*_*u*_) denote *L*_min_(*τ*_*u*_; *N, Z, D, α*).

To study the behavior of *L*_min_(*τ*_*u*_) as *τ*_*u*_ → ∞, we define

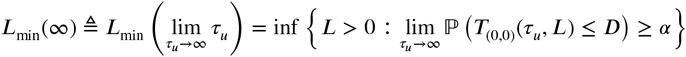

where the last equality holds because the limit and probability can be exchanged for the same reason as in section C. In the limit of *τ*_*u*_ → ∞, the unbinding rate *k*_*u*_ → 0, and therefore the rates *Q*_(*n,z*),(*n*−1,*z*)_ = *Q*_(*n,z*),(*n*−1,*z*−1)_ → 0, i.e. the activation time should be approximated by a Markov chain where the state (*n, z*) is always monotonically increasing. To this end, define the Markov chain *M*^(*f*)^ with only forward transitions i.e. the rate matrix *Q*^(*f*)^ is given by

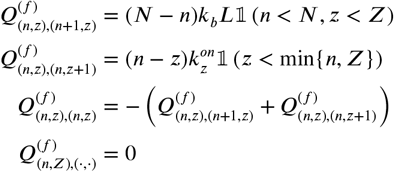

and all other rates not specified are 0. The only difference between the original rate matrix *Q*, and the rate matrix *Q*^(*f*)^ is that 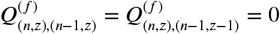. Let 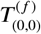 denote the first passage time from the state (0, 0) to the boundary ℬ for the chain *M*^(*f*)^. Then we have the following result.

### Theorem D.1 (Horizontal Asymptote)

*The minimum density L*_min_(∞) *required to activate in the limit τ*_*u*_ → ∞ *is given by*

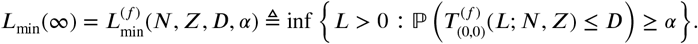

Theorem D.1 implies that we can compute *L*_min_(∞) and understand its dependence on (*N, Z, D, α*) by computing the first passage time distribution for the simpler Markov chain *M*^(*f*)^. Computing the Laplace transforms ℒ _(*i,j*)_(*s*) for *M*^(*f*)^ reduces to solving a sparse upper triangular set of equations. These equations can be solved via back-substitution resulting in 𝒪(*NZ*) complexity as compared to 𝒪(*N*^2^*z*^2^) for computing the Laplace transforms for the first passage times for *M*. Figure D.1 confirms that 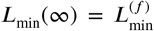 and that 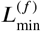 can be computed much faster than *L*_min_(∞) (∼10-100 times faster for moderate-sized *N* and *Z*). Computing the horizontal asymptote is faster for smaller *N* and *z* because it only requires one pass through the problem data (of size 𝒪(*NZ*)), whereas computing the vertical asymptote requires two passes through the problem data (of size 𝒪(*Z*)). The difference in complexity starts to show up once *N* and/or *Z* become larger.

*Proof.* We equivalently show that

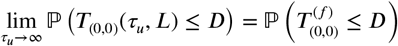

As in the analysis of the vertical asymptote, it suffices to show

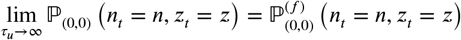

for all (*n, z*) ∈ 𝒮 and *t* > 0, with 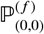 being the distribution of the modified Markov chain *M*^(*f*)^ given (*n*_0_, *z*_0_) = (0, 0). We have

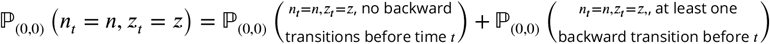

We also see that

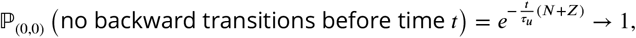

as *τ*_*u*_ → ∞, and

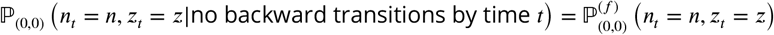

Putting these two together tells us that

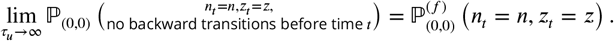

A similar argument establishes that 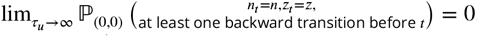. Therefore,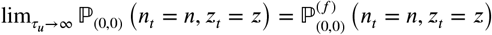. □

## E 1-D approximation for finite *L, τ*_*u*_

In Sections C and D we established that the vertical and horizontal asymptotes of the *L*_*α*_(*τ*_*u*_) curve can be characterized in terms of simpler Markov chains whose first passage time distributions can be computed much more quickly. In this section, we study the shape of the *ℒ*_*α*_ curve for finite values of *L* and *τ*_*u*_. We do this because being able to efficiently calculate *τ*_*α*_(*L*) and ℒ _*α*_ (*τ*) is essential for computing the fidelity metric discussed in section 2.3.

In order to efficiently simulate the entire ℒ _*α*_ curve and investigate its “distance” from ideal absolute ligand distribution behavior, we develop a 1-dimensional approximation of the full model *M* where the *n*- and *z*−dynamics are decoupled. We show that this decoupling significantly reduces the complexity of computing (*τ*_*ϵ*_, *L*_*ϵ*_).

Consider a birth-death process *B* = {*b*(*t*): *t* ≥ 0} on the state space 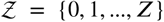 with an absorbing boundary at *Z*. The transition rates are

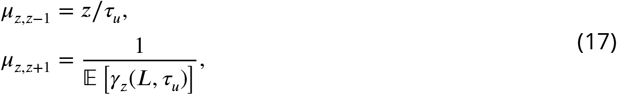

with *γ*_*z*_(*L, τ*_*u*_) being the absorption time in a *z*-dependent Markov chain defined below. See Figure E.1.

**Figure D.1.**
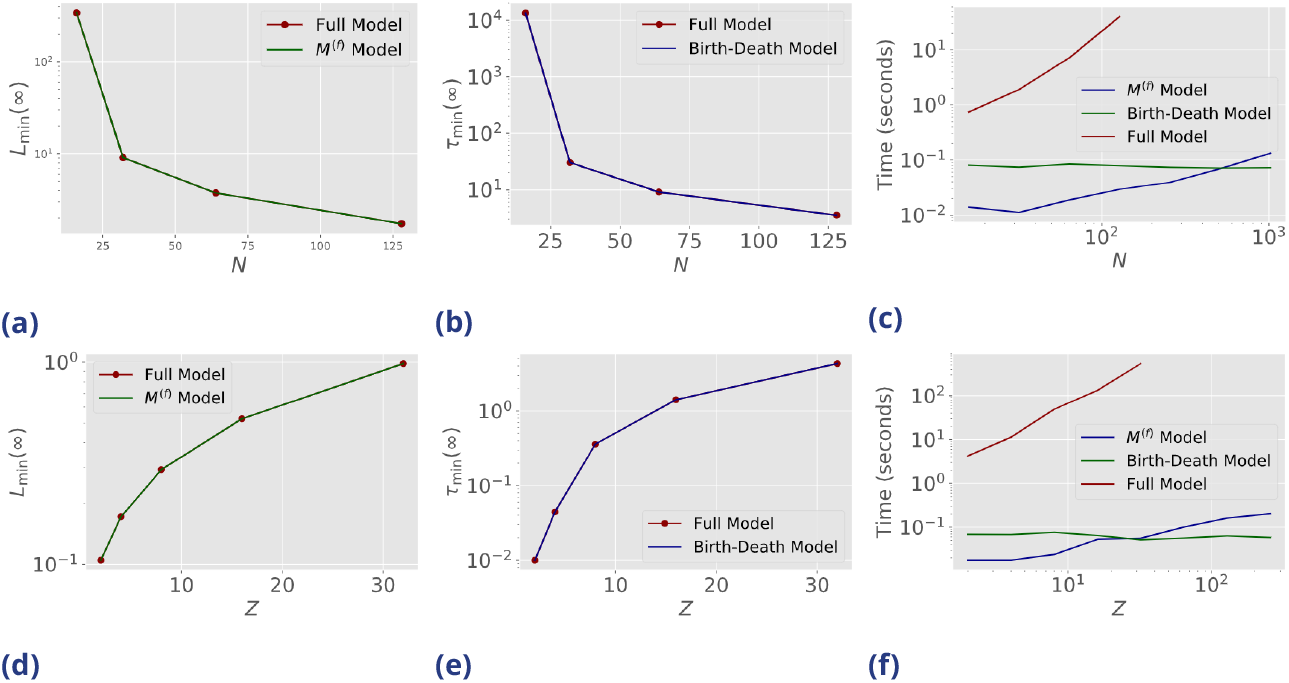
Comparison of location of horizontal and vertical asymptotes computed using the full model vs. using the simplified models, as well as the speed to compute them. (a) *L*_min_(∞) computed using full model (red dots) and simplified model *M*^(*f*)^ (green line), as a function of *N* ∈ {16, 32, 64, 128}, *Z* = 16, *α* = 0.95, *D* = 300s. (b) *τ* _min_(∞) computed using full model and birth-death process (blue line). *N, Z, α, D* same as (a). (c) Speed to compute *L*_min_(∞) using *M*^(*f*)^, speed to compute *τ*_min_(∞) using birth-death process, and speed to compute *τ*_min_(∞) using full model, as a function of *N* (speed to compute *τ*_min_(∞) and speed to compute *L*_min_(∞) using full model are the same). For *L*_min_(∞) and *τ*_min_(∞), *N* values extended to *N* ∈ {16, 32, 64, 128, 256, 512, 1024}. (d)-(f) Same as (a)-(c) but as a function of *Z. N* = 256, *α* = 0.95, *D* = 450s. *Z* ∈ {2, 4, 8, 16, 32} for (a), (b), and full model in (c); *Z* ∈ {2, 4, 8, 16, 32, 64, 128, 256} in (c) for simplified models.

Let *M*^(*z*)^ = {*n*^(*z*)^(*t*) : *t* ≥ 0} denote a Markov chain on {*z, z* + 1, …, *N*} ∪ {𝒜}, where the state 𝒜 is an absorbing state. For state *i* ∈ {*z, z* + 1, …, *N*} the transition rates are given by

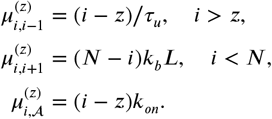

The Markov chain *M*^(*z*)^ starts from the initial state *n*^(*z*)^(0) defined below. Before describing the procedure for selecting *n*^(*z*)^(0), we describe the rationale for this decoupling of the (*n, z*) dynamics.

The T-cell activates when *z* = *Z*. We would be able to efficiently estimate the quantities of interest if one could describe a Markov chain where the transition rates are only a function of the state *z*. The rate from state *z* to state *z* − 1 in the 2-D Markov chain *M* is given by 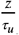, i.e. it only depends on *z*, and so we keep this rate as it is. However, the rate from state *z* to *z* + 1 depends on the number of TCR-antigen complexes *n*, and therefore cannot be written as a function of only the state *z*. We know that *n* ≥ *z*, and so define a Markov chain *M*^(*z*)^ with the state space consisting of {*z*, …, *N*} and an absorbing state A that represents the event *z* → *z* + 1. Note that in the 2-D Markov chain, it’s possible that *n* is much larger than *z* when the chain makes a transition from *z* → *z* + 1, and so setting *n*^(*z*)^(0) = *z* could significantly overestimate the first passage time. To correct this, we set the initial states *n*^(*z*)^(0) in the following recursive manner:

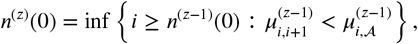

i.e. *n*^(*z*)^(0) is set to the value of *i* ≥ *n*^(*z*−1)^(0) where the rate of transition to the stopping state *𝒜* is larger than the rate to the state *i*+1. Setting *n*^*z*^(0) in this manner allows the approximation to mimic the behaviour of the 2-D chain *M* both for when *Lτ*_*u*_ is very large and the transitions in *M* are likely to be close to (0, 0) → (1, 0) → (2, 0) → (*N*, 0) → (*N*, 1) … → (*N, Z*), and when *Lτ*_*u*_ is very small and the the transitions in *M* are likely to be close to (0, 0) → (1, 0) → (1, 1) → (2, 1) → … → (*N, Z* − 1) → (*N, Z*). This setting also allows for more complicated transitions associated with moderate values of *Lτ*_*u*_. Finally, we replace the first passage time *γ*_*z*_(*L, τ*_*u*_) from *n*_*z*_(0) to 𝒜 by the maximum entropy non-negative random variable with the same mean, i.e. an exponential random variable with rate 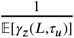. This explains the forward rate in (17).

**Figure E.1.**
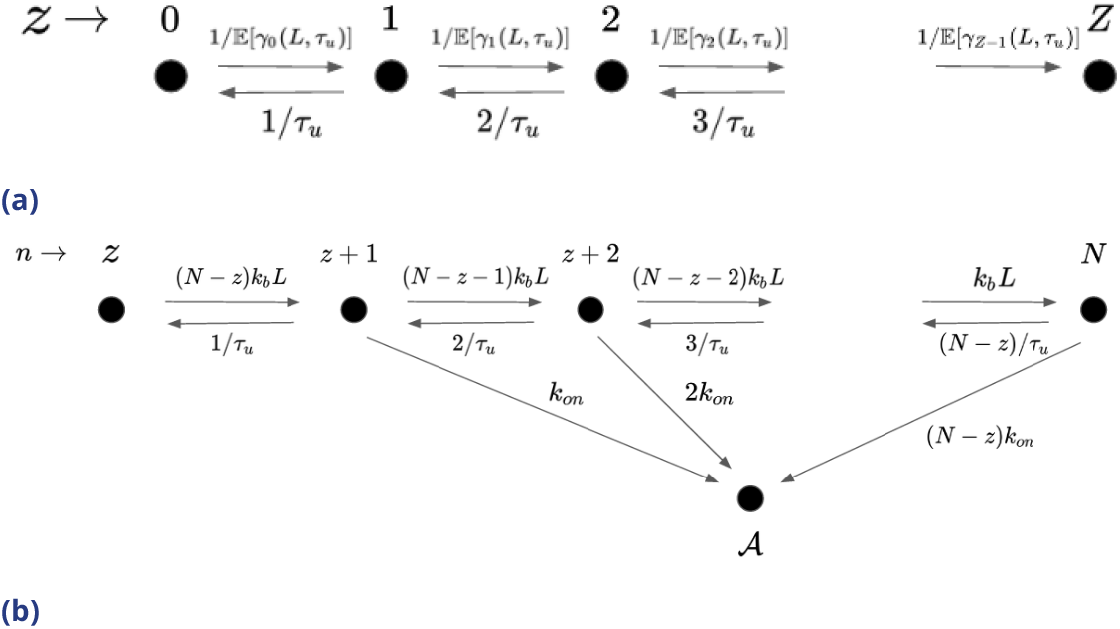
Proposed model used to approximate the convergence of *τ*_min_(*L*) →*τ*_min_(∞) and *L*_min_(*τ*_*u*_) → *L*_min_(∞). The first passage time to the boundary ℬ in *M* is approximated with the first passage time to *z* in the birth-death process shown in (a). *γ*_*z*_(*L, τ*_*u*_) denotes the first passage time to A starting value *n*^(*z*)^(0) in the chain displayed in (b). We replace *γ*_*z*_(*L, τ*_*u*_) by the maximum entropy distribution with mean 𝔼 [*γ*_*z*_(*L, τ*_*u*_)], i.e. an exponential with rate 1/𝔼 [*γ*_*z*_(*L, τ*_*u*_)]. Once the model in subfigure (a) transitions to a new value of *z* (so either to *z* − 1 or *z* + 1), the model here changes to reflect the change in *z* and restarts.

Inverting the Laplace transform for the birth-death process *B* allows us to approximate *τ*_*ϵ*_ and *L*_*ϵ*_. Since calculating the mean first passage time to 𝒜 for each of the *M*^(*z*)^ processes involves solving a tri-diagonal system of equations, this step takes at most 𝒪(*N*) time, resulting in a total complexity of 𝒪(*NZ*). The Laplace transform for the birth-death process can be computed in 𝒪(*Z*) time. Thus, the approximate model results in an 𝒪(*NZ*) speedup compared to using the model *M*.

Figures E.2 and E.3 compare the values of *τ*_*ϵ*_(*N, Z, D, α*) and *L*_*ϵ*_(*N, Z, D, α*) computed using the full model compared to the values computed using the approximation detailed in this section. We see that the true and approximated values of *τ*_*ϵ*_ and *L*_*ϵ*_ are typically within a factor of 2 from one another, and the time to approximate these two values is 10 − 100 times faster for the values of *N* and *Z* tested.

**Figure E.2.**
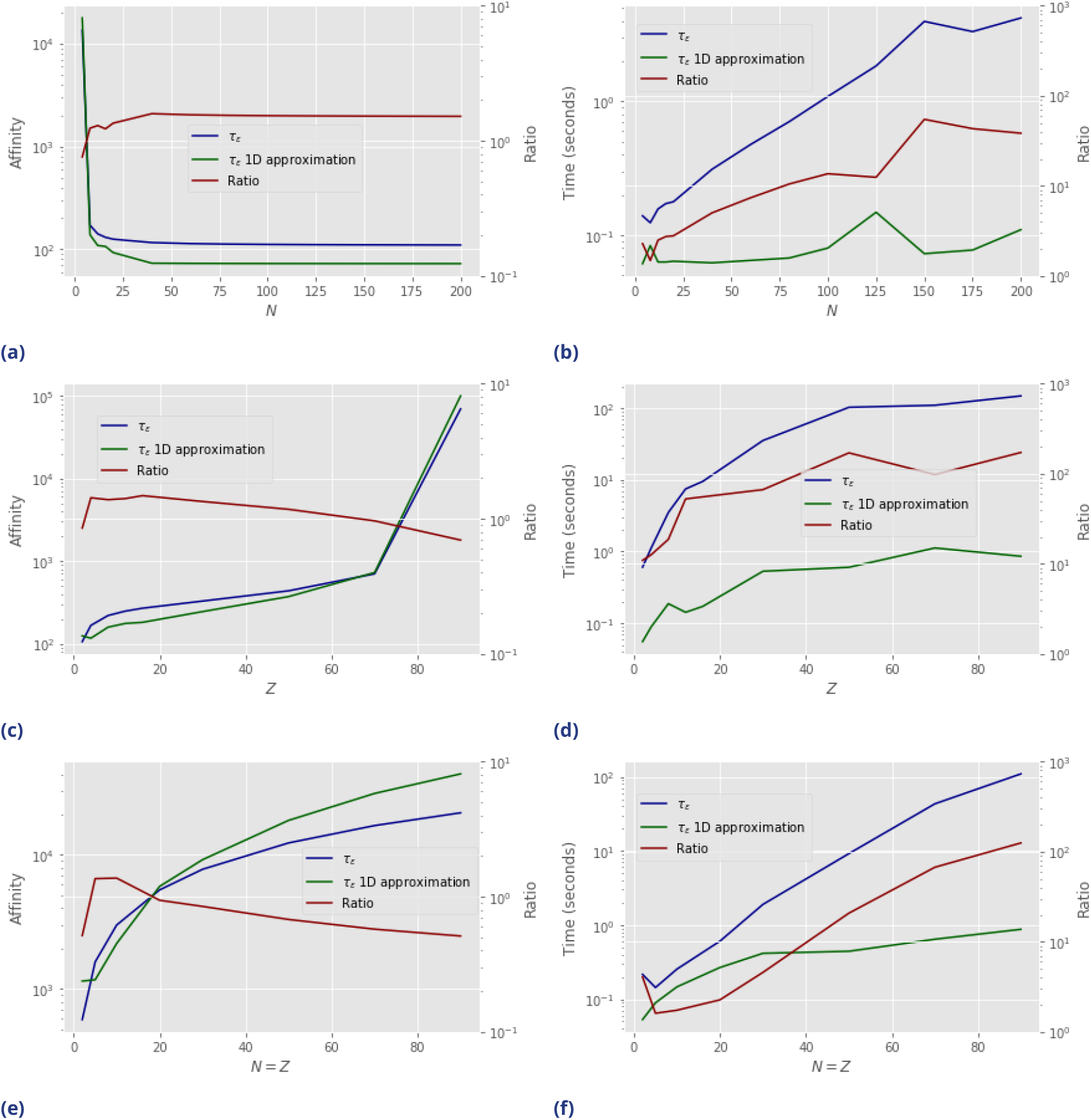
Comparison of *τ*_*ϵ*_ using the full model *M* and the approximation in Figure E.1. *ϵ* = 1 and *α* = 0.95 for all figures. (a) *τ*_*ϵ*_ as computed by the full model *M* (blue line) and the approximation (green line), as well as the ratio of the two (red line), as a function of *N* = 4, …, 200. *Z* = 4, *D* = 225. (b) Time to compute *τ*_*ϵ*_ using original model *M* (blue line) and approximation (blue line), as well as ratio between the two, as a function of *N*. Same parameters as (a). (c) Same as (a) but as a function of *Z* = 2, …, 90. *N* = 90, *D* = 400. (d) Same as (b), but as a function of *Z*. Same parameters as (c). (e) Same as (a), but as a function of *N* = *Z* = 2, …, 90, with *D* set to 1.25*D*^*^(*N, Z, α*). (f) Same as (b), but as a function of *N* = *Z*. Same parameters as (e).

**Figure E.3.**
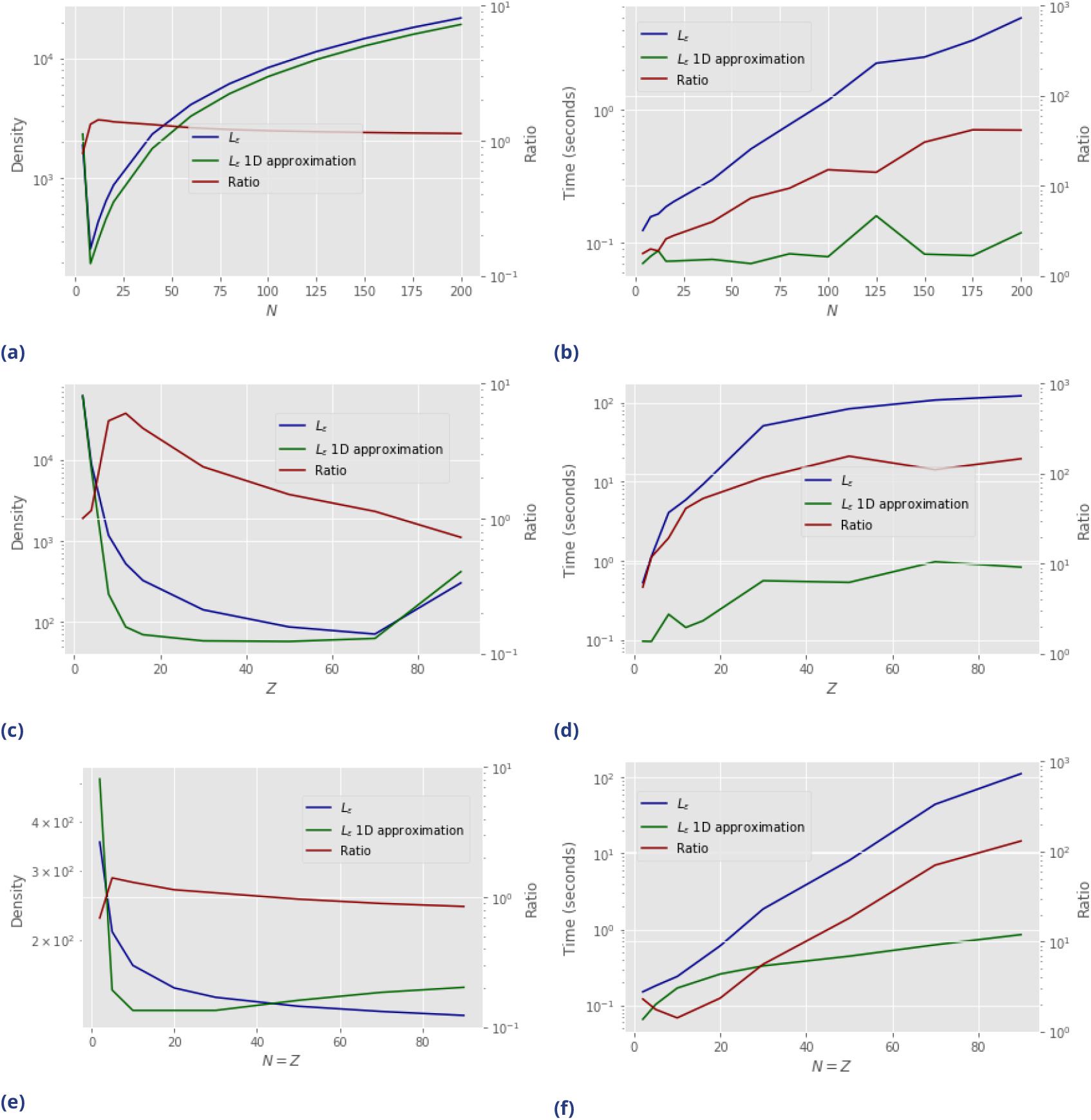
Comparison of *L*_*ϵ*_ using original model *M* and approximation from Figure E.1. Same layout as Figure E.2.

## Footnotes

1 The details of the second molecule, i.e. whether it is phosphorus, Lck, ZAP, etc., are not relevant to our model.

2 For *Z* = 1, the dynamics for ((1 + *ϵ*)*L, τ*_*u*_/(1 + *ϵ*)) are the same as that for (*L, τ*_*u*_). Therefore, absolute ligand discrimination is impossible.

